# The CB_1_ receptor interacts with cereblon and drives cereblon deficiency-associated memory shortfalls

**DOI:** 10.1101/2023.07.24.550332

**Authors:** Carlos Costas-Insua, Alba Hermoso-López, Estefanía Moreno, Carlos Montero-Fernández, Alicia Álvaro-Blázquez, Rebeca Diez-Alarcia, Irene B. Maroto, Paula Morales, Enric I. Canela, Vicent Casadó, Leyre Urigüen, Luigi Bellocchio, Ignacio Rodríguez-Crespo, Manuel Guzmán

## Abstract

Cereblon/CRBN is a substrate-recognition component of the Cullin4A-DDB1-Roc1 E3 ubiquitin ligase complex. Destabilizing mutations in the human *CRBN* gene cause a form of autosomal recessive non-syndromic intellectual disability (ARNSID) that is modelled by knocking-out the mouse *Crbn* gene. A reduction in excitatory neurotransmission has been proposed as an underlying mechanism of the disease, but the intimate factors eliciting this impairment remain mostly unknown. Here we report that CRBN molecules selectively located on glutamatergic neurons are necessary for proper memory function. Combining various *in vivo* approaches, we show that the cannabinoid CB_1_ receptor (CB_1_R), a key suppressor of synaptic transmission, is overactivated in CRBN deficiency-linked ARNSID mouse models, and that the memory deficits observed in these animals can be rescued by acute CB_1_R-selective pharmacological antagonism. Molecular studies demonstrated that CRBN interacts physically with CB_1_R and impairs the CB_1_R-G_i/o_-cAMP-PKA pathway in a ubiquitin ligase-independent manner. Taken together, these findings unveil that CB_1_R overactivation is a driving mechanism of CRBN deficiency-linked ARNSID and anticipate that the blockade of CB_1_R could constitute a new therapy for this orphan disease.

## Introduction

Intellectual disability (ID), defined by an intelligence quotient (IQ) below 70, affects 1-3% of humans worldwide (Schalock *et al*, 2010). Individuals suffering from ID display impaired cognitive and learning abilities, as well as a compromised adaptability to day-to-day life. Among the many different genes that have been linked to ID (Kochinke *et al*, 2016), *CRBN*, the gene encoding the 442 amino-acid protein cereblon/CRBN, was identified 20 years ago in a study searching for gene(s) causing a non-severe form of autosomal recessive non-syndromic intellectual disability (ARNSID) found in American individuals with German roots (Higgins *et al*, 2000, 2004). These individuals bore a single nucleotide substitution (*CRBN*: c.1255C→T), which generates a premature stop codon (R419X), and displayed memory and learning deficits, with IQ values ranging from 50 to 70. Individuals carrying a different *CRBN* missense mutation (*CRBN*: c.1171T→C; C391R), which gives rise to more aggressive clinical symptoms, were subsequently identified in Saudi Arabia (Sheereen *et al*, 2017). Copy number variations in the chromosomal region containing the *CRBN* gene also result in ID (Dijkhuizen *et al*, 2006; Papuc *et al*, 2015). Despite these well-described pathological consequences of *CRBN* mutations and the high abundance of CRBN in the brain (Higgins *et al*, 2010), the neurobiological actions of this protein remain obscure.

Seminal studies identified CRBN as a substrate adaptor of the Cullin4A-DDB1-Roc1 E3 ubiquitin ligase complex (CRL4^CRBN^) and the molecular target of thalidomide, a drug that, when prescribed to pregnant women for sedative and antiemetic purposes, caused severe malformations in thousands of children (Ito *et al*, 2010; Fischer *et al*, 2014). Despite these severe teratogenic effects, thalidomide and related immunomodulatory drugs, such as pomalidomide and lenalidomide, are currently used to treat lupus, lepra and some haematological malignancies (Asatsuma-Okumura *et al*, 2019b). An increasing body of evidence suggests that both the therapeutic and the teratogenic effects of thalidomide arise from modifications in the specificity of CRBN towards its ubiquitination substrates upon drug binding to this protein (Ito *et al*, 2010; Krönke *et al*, 2014, 2015; Matyskiela *et al*, 2018; Asatsuma-Okumura *et al*, 2019a). In contrast, little is known about the physiological actions of CRBN, particularly in the brain, which could provide a mechanistic basis to explain why mutations in this protein impact cognition. Previous reports support that the CRBN^R419X^ mutation destabilizes the protein by enhancing autoubiquitination, thus suggesting that the ARNSID-associated neuropathology could arise from reduced CRBN levels (Xu *et al*, 2013). Consistently, knocking-out the *Crbn* gene in mice impairs learning and memory (Bavley *et al*, 2018; Choi *et al*, 2018). To date, the proposed mechanisms underlying this CRBN deficiency-associated cognitive impairment remain limited to a dysregulation of large conductance Ca^2+^- and voltage-gated potassium channels (BK_Ca_) and an increased activity of AMP-dependent protein kinase (AMPK). These processes could alter synaptic plasticity and reduce excitatory-neuron firing (Liu *et al*, 2014; Bavley *et al*, 2018; Choi *et al*, 2018).

The type-1 cannabinoid receptor (CB_1_R), one of the most abundant G protein-coupled receptors in the mammalian brain, constitutes the primary molecular target of endocannabinoids (anandamide and 2-arachidonoylglycerol) and Δ^9^- tetrahydrocannabinol (THC), the main psychoactive component of the hemp plant *Cannabis sativa* (Pertwee *et al*, 2010). By reducing synaptic activity through heterotrimeric G_i/o_ protein-dependent signalling pathways, the CB_1_R participates in the control of multiple biological processes, such as learning and memory, motor behaviour, fear and anxiety, pain, food intake and energy metabolism (Piomelli, 2003; Mechoulam *et al*, 2014). Specifically, in the context of the present work, cannabinoid-evoked CB_1_R stimulation impairs various short- and long-term cognitive functions in both mice (Figueiredo & Cheer, 2023) and humans (Crean *et al*, 2011; Dellazizzo *et al*, 2022). Given that *Crbn* knockout mice show a reduced excitatory firing and ID-like cognitive impairments, we hypothesized that a pathological CB_1_R overactivation could underlie CRBN deficiency-induced ID. By developing new conditional *Crbn* knockout mouse lines and combining a large number of *in vitro* approaches with extensive *in vivo* behavioural phenotyping, here we show that *i)* the pool of CRBN molecules located on telencephalic glutamatergic neurons is necessary for proper memory function; *ii)* CRBN interacts physically with CB_1_R and inhibits receptor-coupled G_i/o_ protein-mediated signalling; *iii*) CB_1_R is overactivated in CRBN-deficient mice; and *iv)* acute CB_1_R-selective pharmacological blockade rescues the memory deficits induced by genetic inactivation of the *Crbn* gene. These preclinical findings might pave the way to the design of a new therapeutic intervention aimed to treat cognitive symptoms in patients with CRBN deficiency-linked ARNSID.

## Results

### Selective genetic inactivation of *Crbn* in glutamatergic neurons impairs memory

To model *CRBN* mutation-associated ID, we generated three mouse lines in which the *Crbn* gene was selectively inactivated in either *i)* all body cells (hereafter, CRBN-KO mice), *ii)* telencephalic glutamatergic neurons (hereafter, Glu-CRBN-KO mice) or *iii)* forebrain GABAergic neurons (hereafter, GABA-CRBN-KO mice). This was achieved by backcrossing mice carrying exons 3-4 of *Crbn* flanked by *loxP* sites (*Crbn^F/F^*) (Rajadhyaksha *et al*, 2012) with mice expressing Cre recombinase under the control of *i)* the citomegalovirus (*CMV*) promoter, *ii)* the *Nex1* promoter or *iii)* the *Dlx5/6* promoter, respectively (Fig 1A) (Schwenk *et al*, 1995; Monory *et al*, 2006). The three CRBN-deficient mouse lines were viable, fertile, and born at both sexes with expected Mendelian frequency. To evaluate the recombination process together with the neuronal pattern of CRBN expression, we performed *in situ* hybridization experiments in brain sections using RNAscope technology. *Crbn* mRNA was found throughout the brain of CRBN-WT mice, with a remarkable abundance in the hippocampal formation (Fig 1B). High *Crbn* mRNA levels were also detected in the cortex (Fig 1C), striatum (Fig EV1A) and the cerebellum (Fig EV1B). Sections from CRBN-KO mice, as expected, showed a negligible signal in all brain regions analysed (Fig 1B and C, and Fig EV1A and B). In Glu-CRBN-KO mice, *Crbn* mRNA was notably reduced in the CA1, CA3 and hilus of the hippocampus, with a slighter decrease in the granule cell layer of the dentate gyrus (Fig 1B). *Crbn* mRNA was also decreased in the cortex of Glu-CRBN-KO mice (Fig 1C), but not in the striatum (Fig EV1A) and cerebellum (Fig EV1B), two regions that do not express Cre under the *Nex1* promoter (Kleppisch *et al*, 2003). In GABA-CRBN-KO mice, among the four areas analysed, *Crbn* mRNA only diminished in the striatum, a region that is composed almost exclusively by GABAergic neurons (Fig 1B and C, and Fig EV1A and B). All these changes in *Crbn* mRNA levels were confirmed by quantitative PCR (Fig 1D and Fig EV1C) and occurred in concert with changes in CRBN protein levels, as assessed by western blotting (Fig 1E and Fig EV1D). Taken together, these data indicate that CRBN is largely expressed in glutamatergic neurons of the mouse hippocampus and cortex.

**Figure 1.**
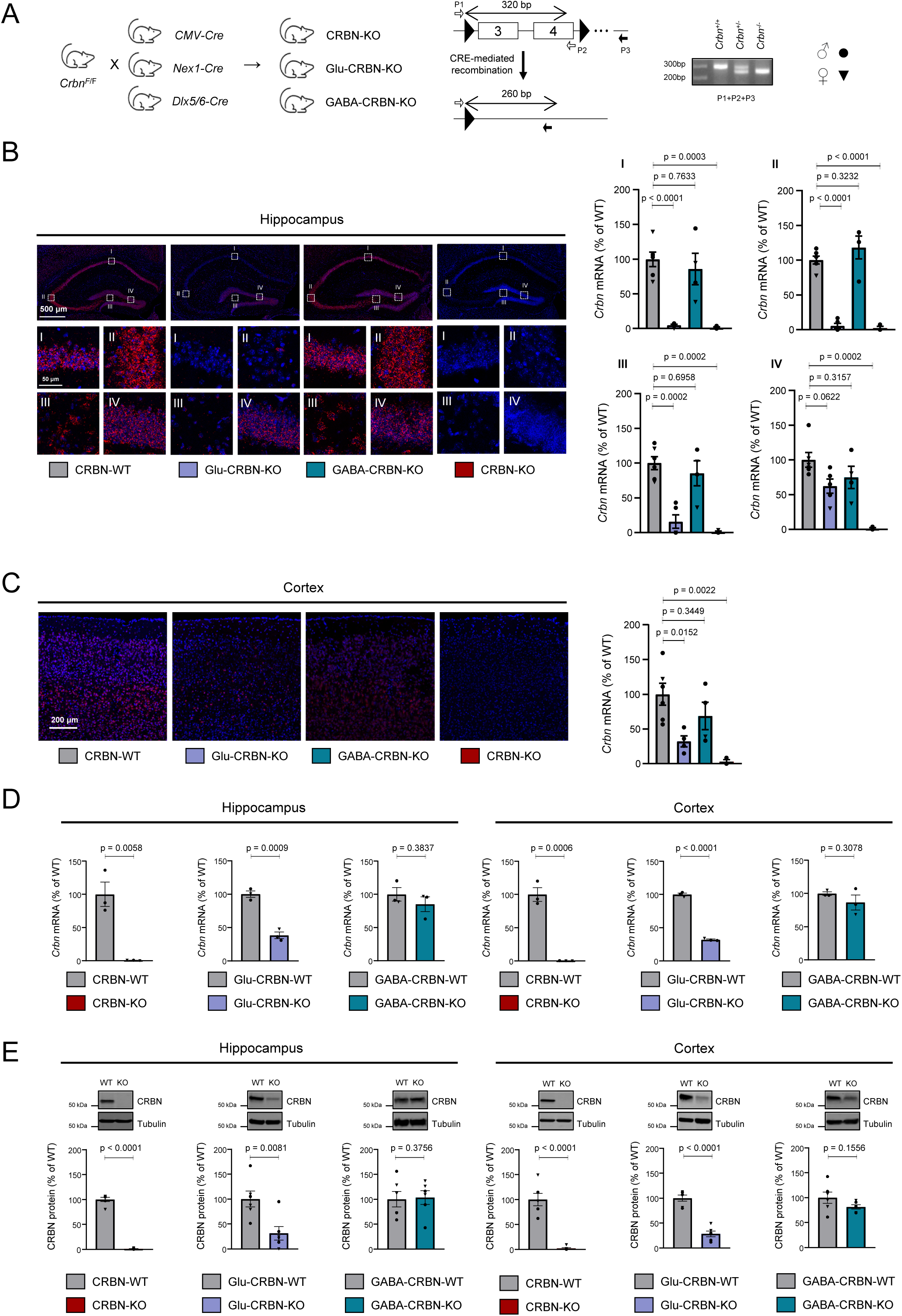
Characterization of the conditional CRBN knockout mouse lines. A. Scheme of the breeding strategy. The resulting genomic architecture, sequencing primers and a representative genotyping agarose gel are shown. B. Representative images and fluorescent signal quantification of RNAscope *in situ* hybridization labelling of *Crbn* mRNA in the hippocampus of CRBN-WT (n = 6), Glu-CRBN-KO (n = 5), GABA-CRBN-KO (n = 4) and CRBN-KO (n = 3) mice. High magnification images of CA1 (I), CA3 (II), hilus (III) and granule cell layer of the dentate gyrus (IV) are shown. Circles, male mice; triangles, female mice. p values were obtained by one-way ANOVA with Dunnett’s post-hoc test. C. Representative images and fluorescent signal quantification of RNAscope *in situ* hybridization labelling of *Crbn* mRNA in the cortex of CRBN-WT (n = 6), Glu-CRBN-KO (n = 4), GABA-CRBN-KO (n = 4) and CRBN-KO (n = 3) mice. Circles, male mice; triangles, female mice. p values were obtained by one-way ANOVA with Dunnett’s post-hoc test. D. *Crbn* mRNA levels (% of WT mice) as assessed by q-PCR in the hippocampus and cortex of CRBN-WT, CRBN-KO, Glu-CRBN-WT, Glu-CRBN-KO, GABA-CRBN-WT and GABA-CRBN-KO mice (n = 3 animals per group). Circles, male mice; triangles, female mice. p values were obtained by unpaired Student’s *t* test. E. CRBN protein levels (% of WT mice) as assessed by western blotting the in hippocampus and cortex of CRBN-WT, CRBN-KO, Glu-CRBN-WT, Glu-CRBN-KO, GABA-CRBN-WT and GABA-CRBN-KO mice (n = 6 animals per group). Circles, male mice; triangles, female mice. p values were obtained by unpaired Student’s *t* test.

Next, we characterized these mice from a behavioural standpoint. CRBN-KO, Glu-CRBN-KO and GABA-CRBN-KO animals showed normal functional parameters such as body weight and body temperature (Fig 2A and B), motor activity (Fig 2C), motor learning (Fig 2D) and gait pattern (Fig EV2A) compared to control CRBN-floxed littermates. Anxiety-like behaviour, as assessed by the elevated plus maze test (Fig 2E) or the number of entries in the central part of an open-field arena (Fig EV2B), was also unchanged between genotypes. As a previous study had linked alterations in *CRBN* copy number to autism spectrum disorders (Pinto *et al*, 2010), we evaluated sociability and depression, two core symptoms of those disorders, using the three-chamber test and the forced-swimming test, respectively. CRBN-KO, Glu-CRBN-KO and GABA-CRBN-KO mice had a preserved sociability (Fig 2F) and did not show major signs of depression (Fig 2G) compared to matched controls. Regarding memory function, which is heavily impaired in individuals bearing CRBN mutations (Higgins *et al*, 2000, 2004), first, we found that long-term recognition memory was compromised in CRBN-KO mice when using the novel object recognition test. Of note, this trait required CRBN molecules located on excitatory neurons, as Glu-CRBN-KO, but not GABA-CRBN-KO, also underperformed in the task (Fig 2H). To further strengthen this notion, we used a modified version of the Y-maze test aimed to evaluate spatial memory. Again, CRBN-KO and Glu-CRBN-KO mice, but not GABA-CRBN-KO, travelled less distance in a novel arm compared to a previously familiar arm, in contrast with their control littermates (Fig 2I). Finally, as an additional memory-related measure, we used a contextual fear-conditioning paradigm. We previously verified that pain sensitivity, using the hot plate test, was not basally affected by knocking-out *Crbn* (Fig EV2C), and that the freezing response was unaltered during the shocking session (Fig EV2D). In line with the aforementioned observations, we found that, compared to CRBN-floxed mice, the aversive stimulus elicited a lower freezing response in CRBN-KO and Glu-CRBN-KO mice, but not in GABA-CRBN-KO animals, when reintroduced in the shocking chamber 24 h after conditioning (Fig 2J). Taken together, these data show that knocking-out *Crbn* in mice, while preserving most behavioural traits, causes a remarkable memory impairment, and underline the necessity for CRBN molecules selectively located on telencephalic excitatory neurons for a proper cognitive function.

**Figure 2.**
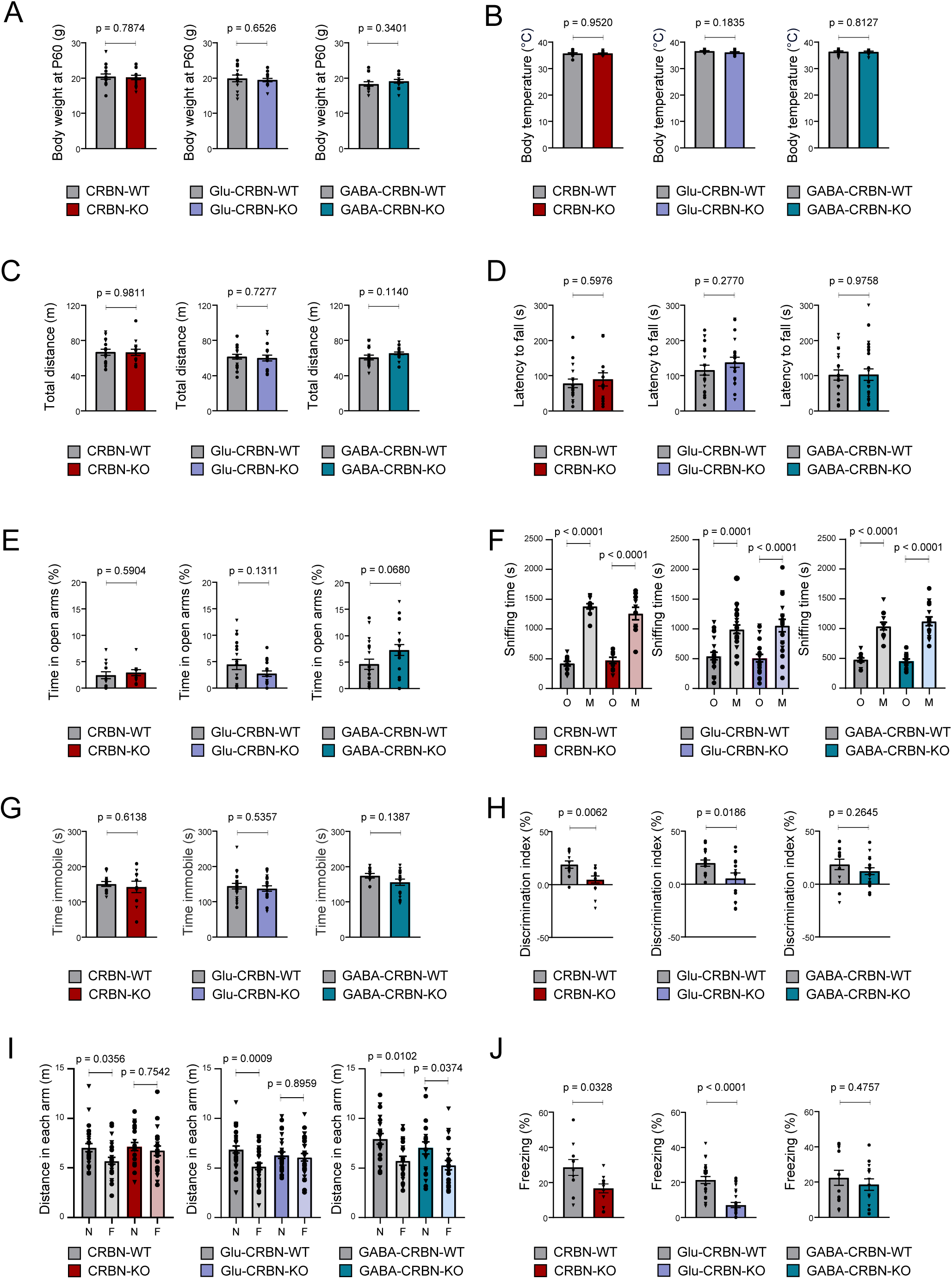
Behavioural phenotyping of the conditional CRBN knockout mouse lines. A. Body weight (in g) at postnatal day 60. CRBN-WT (n = 16), CRBN-KO (n = 16), Glu-CRBN-WT (n = 16), Glu-CRBN-KO (n = 16), GABA-CRBN-WT (n = 16), GABA-CRBN-KO (n = 16). Circles, male mice; triangles, female mice. p values were obtained by unpaired Student’s *t* test. B. Body temperature (in °C) at postnatal day 60. CRBN-WT (n = 12), CRBN-KO (n = 12), Glu-CRBN-WT (n = 13), Glu-CRBN-KO (n = 12), GABA-CRBN-WT (n = 12), GABA-CRBN-KO (n = 12). Circles, male mice; triangles, female mice. p values were obtained by unpaired Student’s *t* test. C. Ambulation (total distance travelled, in m) in the open field test. CRBN-WT (n = 18), CRBN-KO (n = 15), Glu-CRBN-WT (n = 20), Glu-CRBN-KO (n = 19), GABA-CRBN-WT (n = 20), GABA-CRBN-KO (n = 21). Circles, male mice; triangles, female mice. p values were obtained by unpaired Student’s *t* test. D. Time (in s) to fall from the apparatus in the rotarod test. CRBN-WT (n = 18), CRBN-KO (n = 15), Glu-CRBN-WT (n = 22), Glu-CRBN-KO (n = 20), GABA-CRBN-WT (n = 20), GABA-CRBN-KO (n = 24). Circles, male mice; triangles, female mice. p values were obtained by unpaired Student’s *t* test. E. Time (in %) spent in the open arms of an elevated plus maze. CRBN-WT (n = 13), CRBN-KO (n = 11), Glu-CRBN-WT (n = 19), Glu-CRBN-KO (n = 18), GABA-CRBN-WT (n = 19), GABA-CRBN-KO (n = 20). Circles, male mice; triangles, female mice. p values were obtained by unpaired Student’s *t* test. F. Time (in s) spent sniffing the cage containing an object (O) or a mouse counterpart (M) in the sociability test. CRBN-WT (n = 11), CRBN-KO (n = 10), Glu-CRBN-WT (n = 22), Glu-CRBN-KO (n = 20), GABA-CRBN-WT (n = 11), GABA-CRBN-KO (n = 15). Circles, male mice; triangles, female mice. p values were obtained by one-way ANOVA with Tukey’s post-hoc test. G. Time (in s) spent immobile in the forced-swimming test. CRBN-WT (n = 12), CRBN-KO (n = 10), Glu-CRBN-WT (n = 22), Glu-CRBN-KO (n = 20), GABA-CRBN-WT (n = 11), GABA-CRBN-KO (n = 16). Circles, male mice; triangles, female mice. p values were obtained by unpaired Student’s *t* test. H. Discrimination index values (in %) in the novel object recognition test. CRBN-WT (n = 12), CRBN-KO (n = 14), Glu-CRBN-WT (n = 17), Glu-CRBN-KO (n = 15), GABA-CRBN-WT (n = 13), GABA-CRBN-KO (n = 18). Circles, male mice; triangles, female mice. p values were obtained by unpaired Student’s *t* test. I. Ambulation (total distance travelled, in m) in the novel (N) or familiar (F) arm in the Y-maze memory test. CRBN-WT (n = 26), CRBN-KO (n = 21), Glu-CRBN-WT (n = 32), Glu-CRBN-KO (n = 28), GABA-CRBN-WT (n = 20), GABA-CRBN-KO (n = 22). Circles, male mice; triangles, female mice. p values were obtained by one-way ANOVA with Tukey’s post-hoc test. J. Time (in %) spent freezing in the testing session of the fear conditioning protocol. CRBN-WT (n = 10), CRBN-KO (n = 10), Glu-CRBN-WT (n = 24), Glu-CRBN-KO (n = 24), GABA-CRBN-WT (n = 13), GABA-CRBN-KO (n = 14). Circles, male mice; triangles, female mice. p values were obtained by unpaired Student’s *t* test.

### CRBN interacts with CB_1_R *in vitro*

CRBN was identified in a recent proteomic study from our group aimed to find new CB_1_R carboxy-terminal domain (CTD)-interacting proteins (Maroto *et al*, 2023) . As CB_1_R activation, by reducing presynaptic neurotransmitter release, can produce amnesia (Wilson & Nicoll, 2002; Figueiredo & Cheer, 2023), and an impaired excitatory neurotransmission has previously been observed in CRBN-KO mice (Choi *et al*, 2018), here we sought to validate whether CRBN is a *bona fide* binding partner of the receptor, and if so, what the functional consequences of this interaction are. First, we produced recombinant hCRBN and hCB_1_R-CTD, and performed fluorescence polarization-based, protein-protein interaction assays. A well-defined, saturable curve was observed, conceivably due to a direct, high-affinity CRBN-CB_1_R-CTD interaction (Fig 3A). Second, we conducted co-immunoprecipitation experiments in the HEK-293T cell line, which indicated an association of CRBN to CB_1_R (Fig 3B, C). Third, BRET assays with a Rluc-tagged version of CB_1_R and a GFP-fused CRBN chimaera also supported the interaction (Fig 3D). Fourth, PLA experiments in cells expressing tagged versions of both proteins showed overt fluorescence-positive *puncta*, consistent with a protein-protein association (Fig 3E).

**Figure 3.**
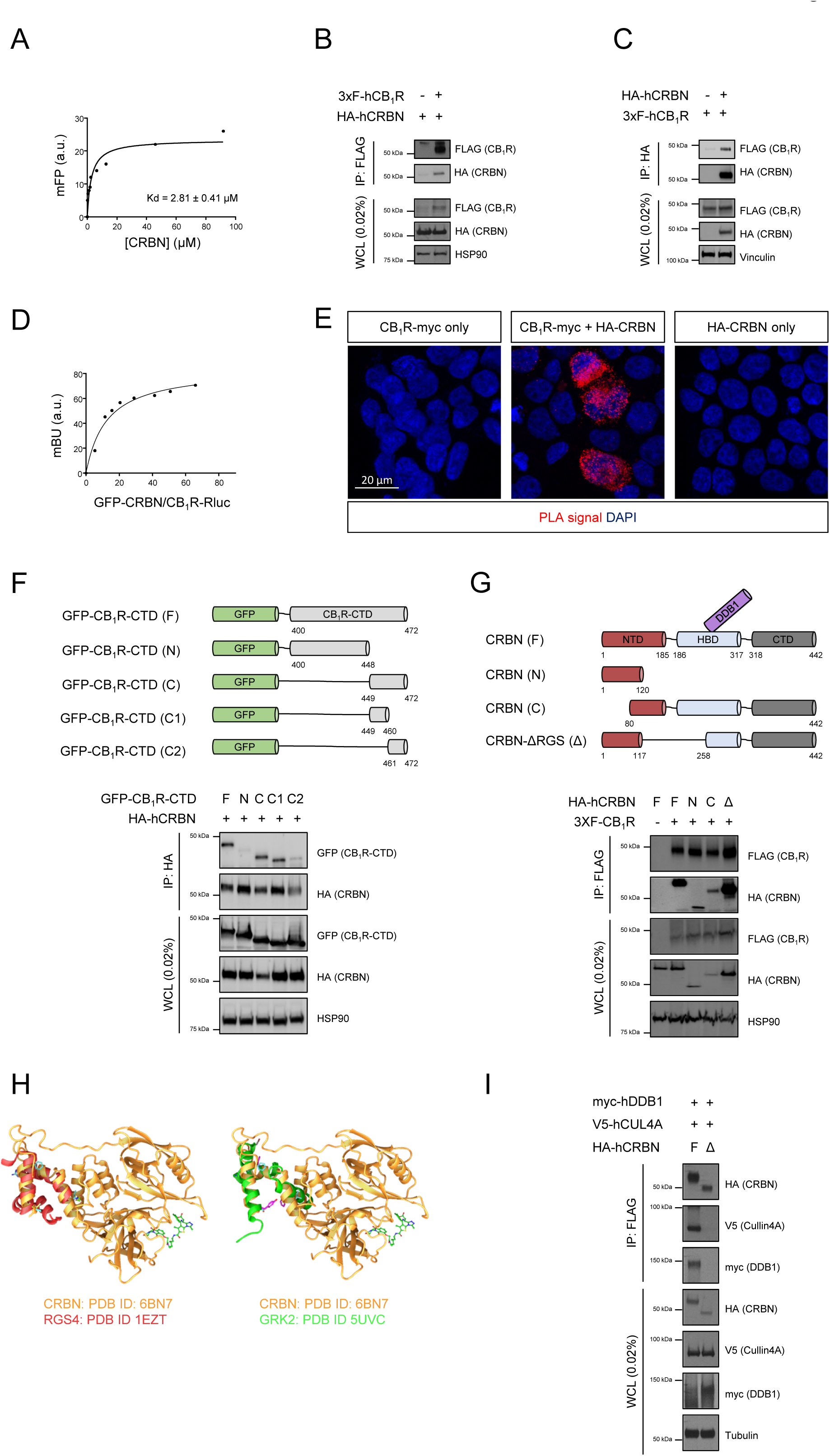
CRBN interacts with CB_1_R *in vitro*. A. Fluorescence polarization-based protein–protein binding experiments using 5-IAF-labeled CB_1_R-CTD and increasing amounts of unlabelled CRBN. A representative experiment is shown (n = 3). B. Co-immunoprecipitation experiments in HEK-293T cells expressing human HA-CRBN and 3xFLAG-CB_1_R. Immunoprecipitation (IP) was conducted with anti-FLAG M2 agarose. WCL, Whole-cell lysate. A representative experiment is shown (n = 3). C. Co-immunoprecipitation experiments in HEK-293T cells expressing human HA-CRBN and 3xFLAG-CB_1_R. Immunoprecipitation (IP) was conducted with anti-HA agarose. WCL, Whole-cell lysate. A representative experiment is shown (n = 3). D. BRET experiments in HEK-293T cells expressing CB_1_R-Rluc and increasing amounts of GFP-CRBN. A representative experiment is shown (n = 3). E. Proximity ligation assays in HEK-293T cells expressing CB_1_R-Rluc, HA-CRBN or both. Note the red *puncta* in the doubly transfected cells. A representative experiment is shown (n = 3). F. Scheme of the different constructs expressing portions of CB_1_R-CTD. Co-immunoprecipitation experiments in HEK-293T cells expressing human HA-CRBN and distinct GFP-CB_1_R-CTD chimeras. Immunoprecipitation (IP) was conducted with anti-HA agarose. WCL, Whole-cell lysate. A representative experiment is shown (n = 3). G. Scheme of the different constructs expressing portions of CRBN. Co-immunoprecipitation experiments in HEK-293T cells expressing human 3xFLAG-CB_1_R and distinct HA-CRBN chimeras. Immunoprecipitation (IP) was conducted with anti-FLAG M2 agarose. WCL, Whole-cell lysate. A representative experiment is shown (n = 3). H. Superposition of the putative RGS domain in CRBN (in gold; Protein Data Bank [PDB] ID: 6BN7) with the RGS domains of RGS4 (left part, in red; Protein Data Bank [PDB] ID: 1EZT) or GRK2 (right part, in green; Protein Data Bank [PDB] ID: 5UVC). Images were constructed with ChimeraX software. I. Co-immunoprecipitation experiments in HEK-293T cells expressing human HA-CRBN (F) or HA-CRBN-ΔRGS (Δ) together with V5-Cullin4A and myc-DDB1. Immunoprecipitation (IP) was conducted with anti-FLAG M2 agarose. WCL, Whole-cell lysate. A representative experiment is shown (n = 3).

Our original proteomic screening was conducted with hCB_1_R-CTD (aa 408-472) (Maroto *et al*, 2023), thus narrowing down *ab initio* the CB_1_R-CRBN binding site to the bulk intracellular, cytoplasm-facing domain of the receptor. Co-immunoprecipitation experiments with several CB_1_R chimaeras (Fig 3F, upper panel) revealed that an 11-amino acid stretch in the mid/distal CB_1_R-CTD (aa 449-460) suffices for CRBN engagement (Fig 3F, lower panel). CRBN has three different domains, namely an *N*-terminal seven-stranded β-sheet, a *C*-terminus containing a cereblon-unique domain that harbours the thalidomide-binding site, and an α-helical bundle linker that is involved in DDB1 binding (Fischer *et al*, 2014) (Fig 3G, upper panel). Unfortunately, we were unable to locate a particular stretch of CRBN that interacts with CB_1_R as both the *N*-terminal and *C*-terminal portions of CRBN bound the receptor (Fig 3G, lower panel). Of note, the existence of a conserved regulator of G protein signalling (RGS) domain spanning amino acids 117-255 (rat protein numbering) of CRBN, which would partially overlap with the CRBN DDB1-binding site, was long proposed (Jo *et al*, 2005). In fact, based on a published CRBN structure (Nowak *et al*, 2018), we aligned this region with the reported RGS domains of RGS4 and GRK2 (Moy *et al*, 2000; Okawa *et al*, 2017) and found a very similar three-dimensional folding (Fig 3H). Hence, we generated a CRBN construct lacking this region (CRBN-ΔRGS) (Fig 3G, upper panel), which was able to bind CB_1_R (Fig 3G, lower panel) and, like similar previously-reported CRBN mutants (*e.g.*, CRBN-ΔMid in Ito *et al*, 2010), did not form the CRL4^CRBN^ complex (Fig 3I). Taken together, these observations support that CB_1_R and CRBN interact through regions encompassing at least an 11-amino acid stretch of the mid/distal CB_1_R-CTD and multiple surfaces of CRBN.

### CRBN inhibits CB_1_R-evoked G_i/o_ protein signalling *in vitro*

To assess whether CRBN binding alters CB_1_R activity, we first conducted dynamic mass redistribution (DMR) assays. We and others have previously used this approach to study global CB_1_R cell signalling (Viñals *et al*, 2015; Costas-Insua *et al*, 2021; Maroto *et al*, 2023). Transfection of HEK-293T cells expressing CB_1_R with a construct encoding CRBN notably reduced the DMR signal evoked by the CB_1_R agonist WIN55,212-2 (WIN) (Fig 4A). Of note, this inhibition was mimicked by CRBN-ΔRGS, thus pointing to a CRL4^CRBN^-independent action. Next, we aimed to dissect which signalling pathways are affected by CRBN. CB_1_R activation inhibits adenylyl cyclase and so reduces intracellular cAMP concentration *via* the α subunit of G_i/o_ proteins (Howlett *et al*, 1986). Using a forskolin-driven cAMP generation assay, we found that both CRBN and CRBN-ΔRGS reduced the ability of CB_1_R to inhibit cAMP production upon activation by its agonists WIN (Fig 4B) and CP-55,940 (CP) (Fig 4C) in a dose-dependent manner. Moreover, this CB_1_R agonist-evoked decrease in cAMP concentration occurred in concert with PKA inactivation, an effect that was also prevented by CRBN (Fig 4D). This action of CRBN on the CB_1_R/cAMP/PKA axis seemed to be pathway-specific, as CB_1_R-triggered ERK activation, another well-characterized receptor signalling pathway (Pertwee *et al*, 2010), was unaffected by CRBN (Fig EV3A). We next evaluated the G protein subtype-coupling profile of CB_1_R in the presence or absence of CRBN or CRBN-ΔRGS. In line with the aforementioned data, CRBN precluded WIN-evoked G_αi1_ and G_αi3_ coupling to CB_1_R, with an apparent slight shift towards G_αo_ engagement (Fig 4E). This effect was evident as well when using HU-210, another CB_1_R agonist (Fig EV3B). CRBN also displaced G_αq/11_ from agonist-engaged CB_1_R (Fig EV3C). The effect of CRBN was largely mimicked by CRBN-ΔRGS (Fig 4E), thus supporting again an independence from the CRL4^CRBN^ complex. As an additional approach, we assessed CB_1_R function in HEK-293T cells in which the *CRBN* gene was knocked-out by CRISPR/Cas9 technology (HEK293T-*CRBN-KO*) (Krönke *et al*, 2015). Compared to the parental *CRBN-WT* cell line, the CB_1_R agonist-evoked reduction of intracellular cAMP concentration was facilitated in *CRBN-KO* cells (Fig 4F), while knocking-out *CRBN* did not affect CB_1_R-mediated ERK activation (Fig EV3D).

**Figure 4.**
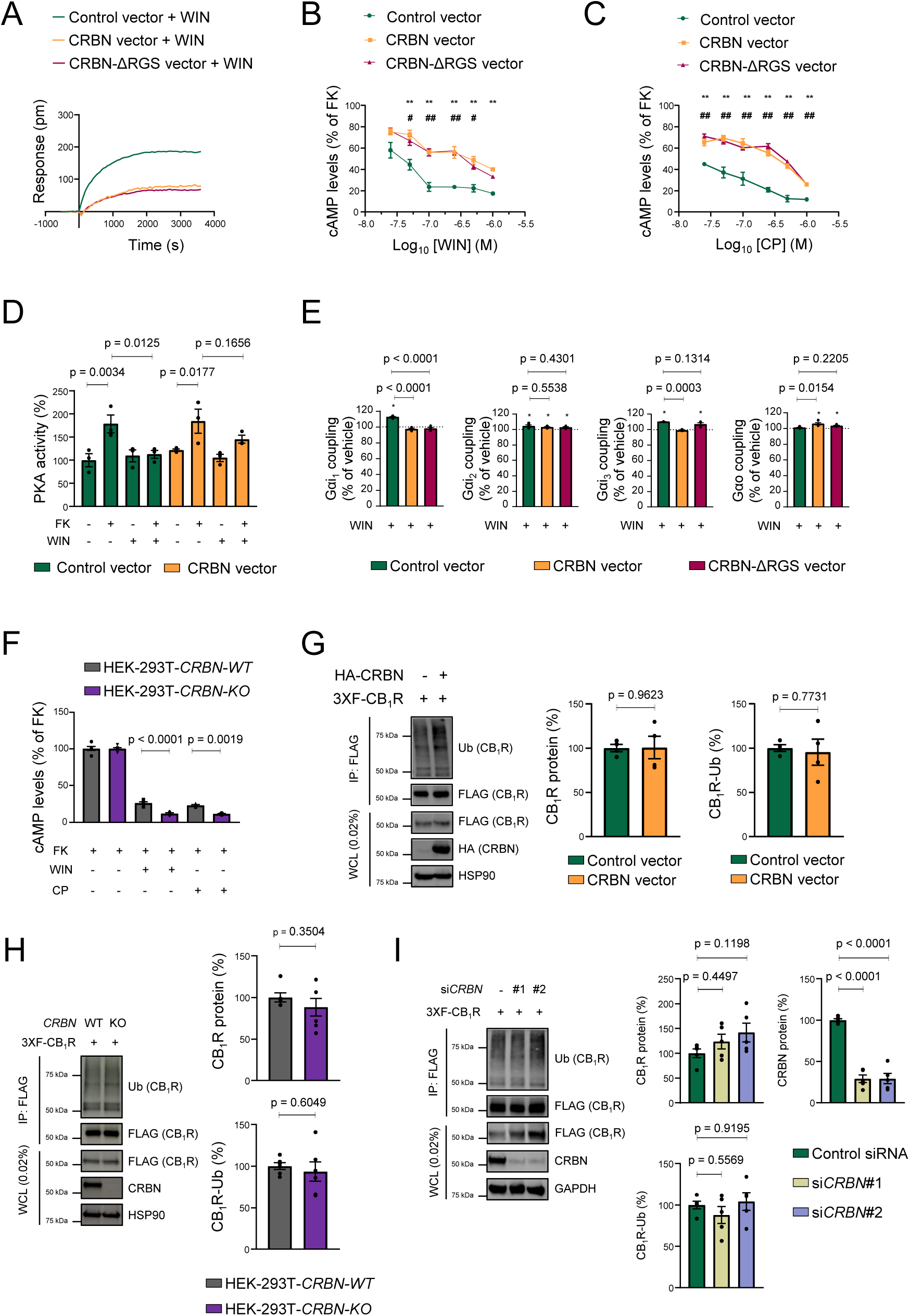
CRBN inhibits CB_1_R-evoked G_i/o_ protein signalling *in vitro*. A. DMR experiments in HEK-293T cells expressing CB_1_R, together or not with CRBN or CRBN-ΔRGS, and incubated with WIN55,212-2 (100 nM). A representative experiment is shown (n = 3). B. cAMP concentration in HEK-293T cells expressing CB_1_R, together or not with CRBN or CRBN-ΔRGS. Cells were incubated first for 15 min with vehicle or WIN55,212-2 (doses ranging from 0.025 to 1 µM), and then for 15 min with forskolin (FK; 500 nM). **p < 0.01 from vehicle, or #p< 0.05 or ##p < 0.01 from paired control, by two-way ANOVA with Tukey’s multiple comparisons test (n = 4). C. cAMP concentration in HEK-293T cells expressing CB_1_R, together or not with CRBN or CRBN-ΔRGS. Cells were incubated first for 15 min with vehicle or CP-55,940 (doses ranging from 0.025 to 1 µM), and then for 15 min with forskolin (FK; 500 nM). p values were obtained by two-way ANOVA with Tukey’s multiple comparisons test (n = 3). D. HEK-293T cells expressing CB_1_R, together or not with CRBN were incubated for 10 min with vehicle or WIN55,212-2 (1 µM) followed by vehicle or forskolin (FK; 1 µM) for another 10 min, and cell extracts were subjected to an ELISA to detect active PKA. Data were normalized to the vehicle-vehicle condition and p values were obtained by two-way ANOVA with Tukey’s multiple comparisons test (n = 3). E. Coupling of CB_1_R to Gα_i/o_ proteins in membrane extracts from HEK-293T cells expressing CB_1_R, together or not with CRBN or CRBN-ΔRGS after WIN55,212-2 stimulation (10 µM). *p<0.05 from basal (dashed line) by one-sample Student’s *t* test. p values between constructs were obtained by unpaired Student’s *t* test (n = 3-4). F. cAMP concentration in HEK-293T-*CRBN-WT* and HEK-293T-*CRBN-KO* cells expressing CB_1_R. Cells were incubated first for 15 min with vehicle, WIN55,212-2 or CP55,940 (each at 500 nM), and then for 15 min with forskolin (FK; 500 nM). p values were obtained by two-way ANOVA with Tukey’s multiple comparisons test (n = 6 for WIN and 3 for CP). G. CB_1_R ubiquitination is not affected by CRBN overexpression. Immunoprecipitation (IP) was conducted with anti-FLAG M2 agarose. WCL: whole-cell lysate. A representative experiment is shown. p values were obtained by unpaired Student’s *t* test (n = 4). H. CB_1_R ubiquitination is not affected by CRBN knockout. Immunoprecipitation (IP) was conducted with anti-FLAG M2 agarose. WCL: whole-cell lysate. A representative experiment is shown. p values were obtained by unpaired Student’s *t* test (n = 6). I. CB_1_R ubiquitination is not affected by CRBN knockdown. Immunoprecipitation (IP) was conducted with anti-FLAG M2 agarose. WCL: whole-cell lysate. A representative experiment is shown. p values were obtained by one-way ANOVA with Tukey’s multiple comparisons test (n = 5).

Aside from these cell-signalling experiments, we evaluated in further detail the possible involvement of ubiquitination as a molecular mechanism by which CRBN could conceivably reduce CB_1_R action. Specifically, we conducted experiments of CRBN *i)* ectopic overexpression (Fig 4G), *ii)* CRISPR/Cas9-based knockout (Fig 4H) and *iii)* siRNA-mediated knockdown (Fig 4I), followed by denaturing immunoprecipitation, and did not find any alteration in CB_1_R levels or ubiquitination. Taken together, these data show that CRBN selectively impairs the CB_1_R-mediated, G_i/o_ protein-coupled inhibition of the cAMP/PKA pathway through a ubiquitination-independent action.

### CRBN interacts with CB_1_R and inhibits receptor signalling in the mouse brain

Our aforementioned *in vitro* experiments support that CRBN binds to and inhibits CB_1_R. Thus, we sought to analyse whether this process also occurs in the mouse brain *in vivo*. As a control, we first verified that the mouse orthologs of CB_1_R and CRBN interact in transfected HEK-293T cells as assessed by co-immunoprecipitation (Fig 5A). We next found that CRBN also co-immunoprecipitates with CB_1_R in mouse hippocampal extracts (Fig 5B). This CB_1_R-CRBN association was further supported by PLA experiments conducted in mouse hippocampal sections, which showed abundant fluorescence-positive *puncta* in WT mice but not CB_1_R-KO animals (Fig 5C). We subsequently injected stereotaxically the hippocampi of WT mice with adenoviral particles encoding a scrambled DNA sequence (AAV1/2.CBA-Control) or FLAG-tagged CRBN (AAV1/2.CBA-FLAG-CRBN) and analysed the G protein-coupling profile of CB_1_R. In line with our aforementioned *in vitro* data, CRBN overexpression occluded the agonist-evoked coupling of CB_1_R to G_αi1_ and G_αi3_ proteins (Fig 5D).

**Figure 5.**
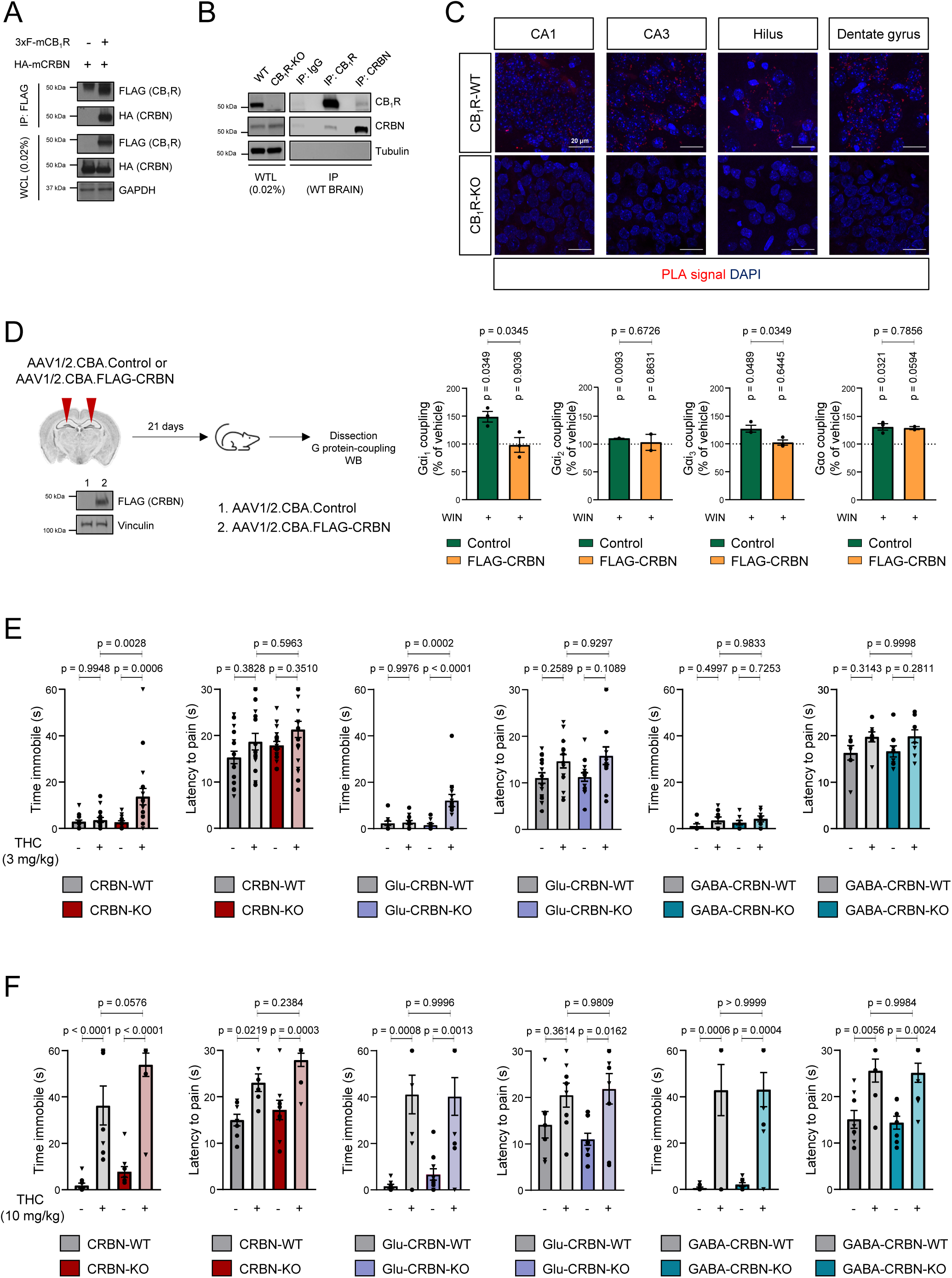
CRBN binds to CB_1_R and inhibits receptor signalling in the mouse brain. A. Co-immunoprecipitation experiments in HEK-293T cells expressing mouse HA-CRBN and 3xFLAG-CB_1_R. Immunoprecipitation (IP) was conducted with anti-FLAG M2 agarose. WCL, Whole-cell lysate. A representative experiment is shown (n = 3). B. Co-immunoprecipitation experiments in adult hippocampal tissue. Immunoprecipitation (IP) was conducted with IgG control, anti-CB_1_R or anti-CRBN. WTL, Whole-tissue lysate. A representative experiment is shown (n = 3). C. Proximity ligation assays in brain slices from WT and CB_1_R-KO mice. Note the fluorescence-positive red *puncta*, depicting CB_1_R-CRBN complexes, in the hippocampus of WT but not KO mice. Representative high magnification images of cortex, CA1, CA3, hilus and granule cell layer of the dentate gyrus are shown (n = 3 animals per group). D. Coupling of CB_1_R to Gα_i/o_ proteins in membrane extracts from hippocampi of mice transduced with AAV1/2.CBA.Control or AAV1/2.CBA.CRBN vectors. p values were obtained by unpaired Student’s *t* test (n = 3) between samples and by one-sample Student’s *t* test from baseline (dashed line). A representative western blot showing viral expression in pooled hippocampal extracts is shown. E. CRBN-WT (n = 16-17), CRBN-KO (n = 18), Glu-CRBN-WT (n = 14-15), Glu-CRBN-KO (n = 14), GABA-CRBN-WT (n = 7-8) and GABA-CRBN-KO (n = 9) mice were injected with a submaximal dose of THC (3 mg/kg, single i.p. injection) or vehicle. Forty min later, catalepsy on a horizontal bar (latency to move, s) and thermal analgesia in the hot-plate test (latency to pain, s) were measured. Circles, male mice; triangles, female mice. p values were obtained by two-way ANOVA with Tukey’s post-hoc test. F. CRBN-WT (n = 6-9), CRBN-KO (n = 9), Glu-CRBN-WT (n = 7-8), Glu-CRBN-KO (n = 9-10), GABA-CRBN-WT (n = 7-8) and GABA-CRBN-KO (n = 8-9) mice were injected with a maximal dose of THC (10 mg/kg, single i.p. injection) or vehicle. Forty min later, catalepsy on a horizontal bar (latency to move, s) and thermal analgesia in the hot-plate test (latency to pain, s) were measured. Circles, male mice; triangles, female mice. p values were obtained by two-way ANOVA with Tukey’s post-hoc test.

CB_1_R activation elicits numerous behavioural alterations in mice, which allows a straightforward procedure to evaluate the status of CB_1_R functionality *in vivo*. Hence, we treated CRBN-deficient mice and their control littermates with vehicle or THC, and assessed two well-characterised cannabinoid-mediated effects, namely catalepsy, which relies exclusively on CB_1_Rs located at CNS neurons (Monory *et al*, 2007), and - as a control-thermal analgesia, which relies mostly on peripherally-located CB_1_Rs (Agarwal *et al*, 2007). Of note, the cataleptic -but not the analgesic-effect induced by a submaximal dose of THC (3 mg/kg) was notably augmented in both CRBN-KO and Glu-CRBN-KO mice, but not in GABA-CRBN-KO mice (Fig 5E). In contrast, a maximal dose of THC (10 mg/kg) induced the same “ceiling” effect in the three mouse lines (Fig 5F), thus supporting a facilitation of CB_1_R function rather than an alteration of global CB_1_R availability. Accordingly, the total levels of hippocampal CB_1_R were not affected upon knocking-out *Crbn* (Fig EV4A, B). The expression of archetypical synaptic markers (vGAT, vGLUT1, synaptophysin, PSD-95) was neither altered in the hippocampi of the three mouse lines compared to matched WT control animals (Fig EV4C). Taken together, these data support that CRBN interacts with CB_1_R and inhibits receptor action *in vivo*.

### Selective pharmacological blockade of CB_1_R rescues CRBN deficiency-associated memory impairment in mice

Finally, we asked whether blocking the aforementioned CB_1_R disinhibition that occurs in CRBN-KO mice could exert a therapeutic effect on these animals by ameliorating their memory deficits. To test this possibility, we treated CRBN-KO mice with a low dose (0.3 mg/kg, single i.p. injection) of the CB_1_R-selective antagonist rimonabant (aka SR141716) prior to behavioural testing. Knocking-out *Crbn* impaired object-recognition memory (Fig 6A, left histogram), freezing behaviour (Fig 6B, left histogram) and spatial memory (Fig 6C, upper histogram) in vehicle-treated mice, and all these severe alterations were effectively rescued by acute rimonabant administration without affecting the basal performance of control CRBN-WT littermates. Of note, this therapeutic effect of rimonabant administration on cognitive traits was also evident in Glu-CRBN-KO mice (Fig 6A, right histogram; B, right histogram; and C, lower histogram). Collectively, these observations are consistent with our cell-signalling and animal-behaviour data, and unveil a therapeutic effect of CB_1_R-selective antagonism on CRBN deficiency-associated memory deficits.

**Figure 6.**
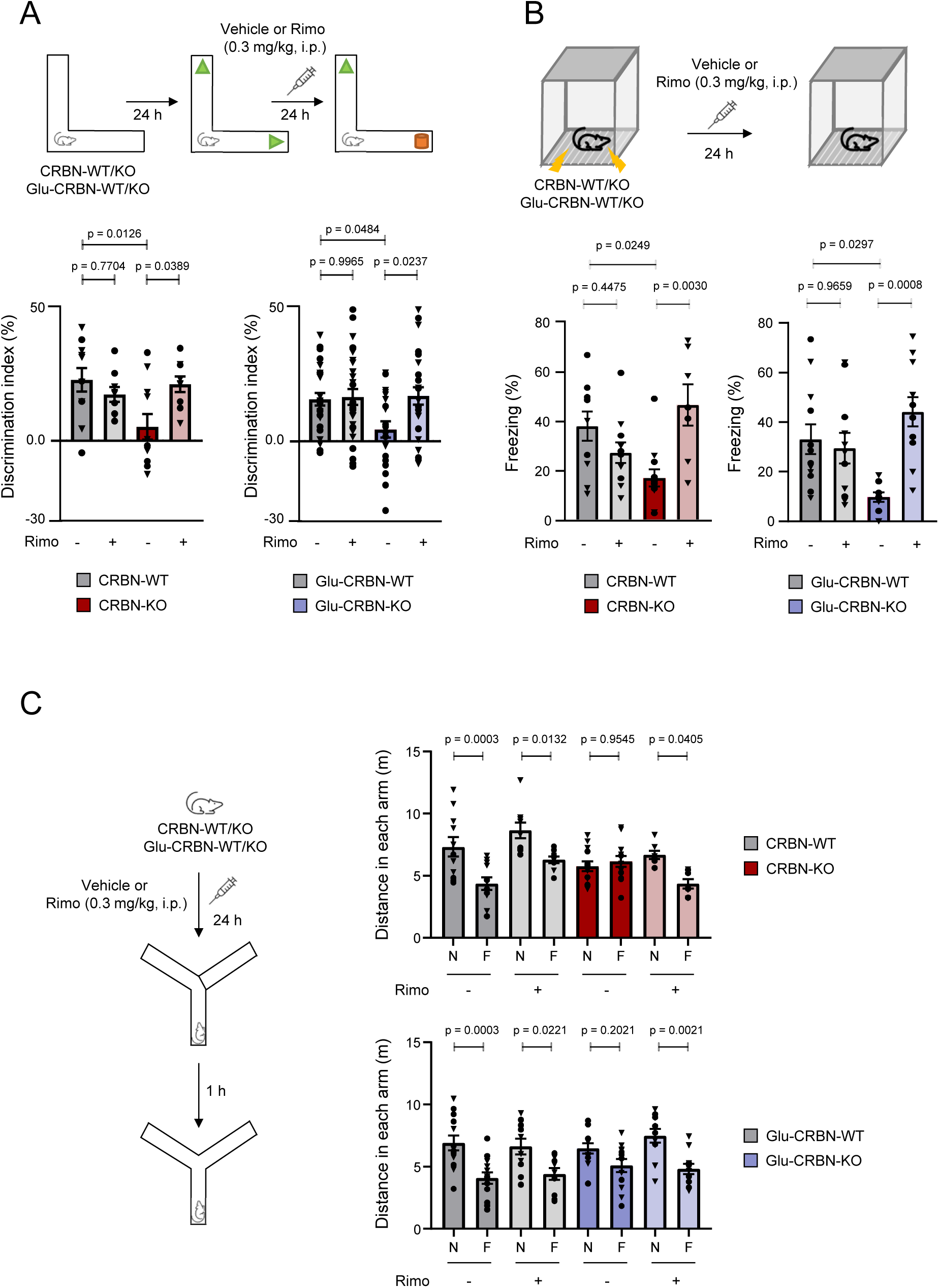
Selective pharmacological blockade of CB_1_R rescues CRBN deficiency-associated memory impairment in mice. A. Experimental scheme and discrimination index values (in %) in the novel object recognition test. CRBN-WT+Veh (n = 11), CRBN-WT+Rimo (n = 9), CRBN-KO+Veh (n = 11), CRBN-KO+Rimo (n = 9), Glu-CRBN-WT+Veh (n = 26), Glu-CRBN-WT+Rimo (n = 28), Glu-CRBN-KO+Veh (n = 21), Glu-CRBN-KO+Rimo (n = 25). Circles, male mice; triangles, female mice. p values were obtained by two-way ANOVA with Tukey’s post-hoc test. B. Experimental scheme and time (in %) spent freezing in the testing session of the fear conditioning protocol. CRBN-WT+Veh (n = 10), CRBN-WT+Rimo (n = 11), CRBN-KO+Veh (n = 12), CRBN-KO+Rimo (n = 7), Glu-CRBN-WT+Veh (n = 12), Glu-CRBN-WT+Rimo (n = 11), Glu-CRBN-KO+Veh (n = 9), Glu-CRBN-KO+Rimo (n = 11). Circles, male mice; triangles, female mice. p values were obtained by two-way ANOVA with Tukey’s post-hoc test. C. Experimental scheme and ambulation (total distance travelled, in m) in the novel (N) or familiar (F) arm in the Y-maze memory test. CRBN-WT+Veh (n = 11), CRBN-WT+Rimo (n = 9), CRBN-KO+Veh (n = 13), CRBN-KO+Rimo (n = 7), Glu-CRBN-WT+Veh (n = 13), Glu-CRBN-WT+Rimo (n = 10), Glu-CRBN-KO+Veh (n = 12), Glu-CRBN-KO+Rimo (n = 11). Circles, male mice; triangles, female mice. p values were obtained by two-way ANOVA with Sidak’s post-hoc test.

## Discussion

Here, upon developing new mouse models lacking CRBN exclusively in telencephalic glutamatergic neurons or forebrain GABAergic neurons, we depicted the neuron-population selectivity of CRBN action. Our mapping of CRBN mRNA and protein expression in the mouse brain shows an enriched expression of CRBN in glutamatergic neurons of the hippocampus, a pivotal area for cognitive performance (Preston & Eichenbaum, 2013). Likewise, our behavioural characterization of those animals demonstrates that Glu-CRBN-KO mice, but not GABA-CRBN-KO animals, display memory alterations. Collectively, this evidence strongly supports that CRBN molecules expressed in hippocampal glutamatergic neurons are necessary for proper memory function, in line with a previous study showing that acute deletion of CRBN from the hippocampus of CRBN-floxed mice (though using a constitutive promoter-driven Cre-recombinase expressing vector) impairs memory traits (Bavley *et al*, 2018). Additional previous work had reported alterations of excitatory neurotransmission in *Crbn* knockout mice (Choi *et al*, 2018). Specifically, an augmented anterograde trafficking and activity of BK_Ca_ channels was suggested to be involved in the reduction of presynaptic neurotransmitter release observed in those animals (Liu *et al*, 2014; Choi *et al*, 2018). Nonetheless, this notion is challenged by other data showing that activation of presynaptic BK_Ca_ channels does not modulate the release of glutamate at several synapses (Gonzalez-Hernandez *et al*, 2018). Our findings may therefore help to reconcile these inconsistencies as CB_1_Rs reduce glutamate release (Piomelli, 2003) and may also activate BK_Ca_ channels under certain conditions (Stumpff *et al*, 2005; Romano & Lograno, 2006; López-Dyck *et al*, 2017). Furthermore, CRBN-KO mice show a resilient phenotype towards stress (Akber *et al*, 2022; Park *et al*, 2022). and the pathological aggregation of Tau, a hallmark of tauopathies as Alzheimer’s disease (Akber *et al*, 2021). Facilitation of CB_1_R signalling also protects against acute and chronic stress, and chronic stress consistently downregulates CB_1_R (Morena *et al*, 2016). A similar scenario occurs in Alzheimer’s disease mouse models, in which CB_1_R pharmacological activation produces a therapeutic benefit and CB_1_R genetic deletion worsens the disease (Aso *et al*, 2012, 2018). Based on our findings, one could speculate that the reported resiliency of CRBN-KO mice may arise, at least in part, from an enhanced CB_1_R-evoked protective activity.

Our array of binding experiments proved that CRBN interacts physically with CB_1_R-CTD, thus highlighting this domain as a molecular hub that most likely influences receptor function in a cell population-selective manner by engaging distinct sets of interacting proteins (Niehaus *et al*, 2007; Costas-Insua *et al*, 2021; Maroto *et al*, 2023). In line with this idea, association with CRBN blunted the ability of CB_1_R to couple to its canonical G_i/o_ protein-evoked inhibition of the cAMP-PKA pathway without altering the receptor ubiquitination status. This effect of CRBN adds to its known ubiquitin ligase-independent, “chaperone-like” actions in the maturation of some membrane proteins (Eichner *et al*, 2016; Heider *et al*, 2021). By doing so, CRBN counteracts the activity of activator of 90-kDa heat shock protein ATPase homolog 1 (AHA1), thereby attenuating its negative effect on membrane protein instability. Intriguingly, chronic CB_1_R activation increases AHA1 levels, and AHA1 has been reported to augment the CB_1_R-mediated effects on cAMP levels and ERK phosphorylation (Filipeanu *et al*, 2011). Therefore, a plausible notion to be explored in the future would be that CB_1_R overactivity upon CRBN loss of function arises, at least in part, from an enhanced, stimulatory action of AHA1 on the receptor.

From a therapeutic perspective, we report that acute CB_1_R-selective pharmacological antagonism fully rescues the memory deficits of both CRBN-KO and Glu-CRBN-KO mice. This finding aligns with previous studies by Ozaita and coworkers, who found improvements in the symptomatology of mouse models of fragile X and Down syndromes upon CB_1_R blockade (Busquets-Garcia *et al*, 2013; Navarro-Romero *et al*, 2019). Rimonabant (Acomplia®) was marketed in Europe for the treatment of obesity until 2008, when it was withdrawn by the EMA due to its severe psychiatric side-effects (Pacher & Kunos, 2013). Of note, the dose of rimonabant used in our study (0.3 mg/kg), when considering a standard inter-species dose conversion formula (Reagan-Shaw *et al*, 2008), is approximately 12 times lower than that prescribed to obesity patients (20 mg/day, equivalent to 3.5 mg/kg in mice), and falls well below the doses reducing food intake (1 mg/kg) and eliciting anxiety (3 mg/kg) in mice (Wiley *et al*, 2005; Thiemann *et al*, 2009). This would theoretically ensure a safer profile upon administration to patients. Given that rimonabant rescues glutamatergic synaptic alterations even at lower doses (0.1 mg/kg) (Gomis-González *et al*, 2016), it is plausible that the dose of 0.3 mg/kg used here normalizes the functionality of the hippocampal circuitry of CRBN-KO and Glu-CRBN-KO mice. These issues notwithstanding, the advent of novel CB_1_R-targeting drugs with a safer pharmacological profile, such as neutral antagonists (*e.g.*, NESS0327) (Meye *et al*, 2013) or negative allosteric modulators (*e.g.*, AEF0117) (Haney *et al*, 2023), constitutes an attractive therapeutic option to be explored in the future.

In summary, we provide compelling evidence supporting the existence of a CRBN-CB_1_R-memory axis that is impaired in *Crbn* knockout mice, thus suggesting that it could also be disrupted in patients with *CRBN* mutations. This study allows a new conceptual view of how CRBN controls memory and provides a potential therapeutic intervention (namely, the pharmacological blockade of CB_1_R) for patients with CRBN deficiency-linked ARNSID. Future work should define the actual translationality of our preclinical-research findings.

## Materials and Methods

### Animals

All the experimental procedures used were performed in accordance with the guidelines and with the approval of the Animal Welfare Committee of Universidad Complutense de Madrid and Comunidad de Madrid, and in accordance with the directives of the European Commission. *Crbn-*floxed mice (herein referred to as *Crbn*^F/F^) and *CMV*-Cre mice were purchased from The Jackson Laboratory (Bar Harbor, ME, USA; #017564, #006054). We also used *Nex1*-Cre mice, *Dlx5/6*-Cre mice and full CB_1_R knockout mice (herein referred to as CB_1_R-KO) (Marsicano *et al*, 2002; Monory *et al*, 2006), which were already available in our laboratory. Animal housing, handling and assignment to the different experimental groups were conducted as described (Ruiz-Calvo *et al*, 2018). Adequate measures were taken to minimize pain and discomfort of the animals. For behavioural experiments, adult mice (*ca.* 2–4-month-old) of both sexes (differentially represented in each graph as circles or triangles) were habituated to the experimenter and the experimental room for one week prior to the experiment. All behavioural tests were conducted during the early light phase under dim illumination (< 50 luxes in the centre of the corresponding maze) and video-recorded to allow the analysis to be conducted by an independent trained experimenter, who remained blind towards the genotype and the treatment of the animal. Mice were weighted on a conventional scale (accuracy up to 0.01 g) and their body temperature was measured with a rectal probe (RET-3, Physitemp, Clifton, NJ, USA) inserted ∼2 cm into the animal’s rectum.

### Motor performance tests

Spontaneous locomotor activity was measured in an open field arena of 70×70 cm built in-house with grey plexiglass. Mice were placed in the centre of the arena and allowed free exploration for 10 min. Total distance travelled, resting time and entries in the central part of the arena (25 × 25 cm) were obtained using Smart3.0 software (Panlab, Barcelona, Spain). To assess motor learning skills, we conducted an accelerating rotarod paradigm consisting of three daily sessions with a 40-min inter-trial interval, for three consecutive days. Briefly, the mouse was placed in the rod (Panlab #LE8205) at a constant speed (4 rpm), which was then accelerated (4 to 40 rpm in 300 s) once the mouse was put in place. The time to fall from the apparatus was annotated in either test, and the mean of trials 4-9 (days 2 and 3) was calculated to ensure reduced inter-trial variability. For gait analysis, mice fore- and hind paws were painted with non-toxic ink of different colours and placed in one end of a corridor (50-cm long, 5-cm wide) on top of filter paper. The distance between strides was measured using a ruler.

### Pain sensitivity test

Analgesia was evaluated using a hot-plate apparatus (Harvard apparatus, Holliston, MA, USA #PY2 52-8570) being the temperature set at 52 °C. Animals were placed in the plate inside a transparent cylinder and latency to first pain symptom (paw licking) was annotated. Mice were removed after 30 s if no symptoms were visible.

### Anxiety test

To evaluate anxiety-like behaviours we employed an elevated plus maze following standard guidelines (arms: 30-cm long, 5-cm wide, two of them with 16-cm high walls, connected with a central structure of 5×5 cm and elevated 50 cm from the floor). Each mouse was placed in the centre of the maze, facing one of the open arms and the exploratory behaviour of the animal was video recorded for 5 min. The number and duration of entries was measured separately for the open arms and the closed arms using Smart3.0 software, being one arm entry registered when the animal had placed both forepaws in the arm. For simplicity, only time of permanence (in %) in the open arms is provided.

### Sociability test

To evaluate social behaviours, we introduced a single mouse in an arena (60-cm long, 40-cm wide, 40-cm high walls) divided in three compartments (20-cm long each) separated by 2 walls (15-cm long) with a connector corridor (10-cm wide) and containing two cylindrical cages (15-cm high, 8.5-cm diameter) in the lateral compartments; for 10 min and allowed free exploration. One h later, the mouse was re-exposed to this environment, but this time one of the cages contained one unfamiliar mouse, paired in sex and age, and being a control genotype with the mouse undergoing testing, in one of the cages. Mouse behaviour was video recorded for 10 min. Finally, time spent sniffing each cage was annotated manually by a blind experimenter using a chronometer. Position of cages containing mice was randomized. Mice with total exploration times lower than 15 s were considered outliers.

### Forced swimming test

The forced swimming test was conducted in a custom square tank (14-cm high, 22-cm wide) filled with 10-cm of water kept at a constant temperature of 22 °C for 5 min. Animal behaviour was video recorded, and time spent immobile was annotated manually by a blind experimenter using a chronometer.

### Novel object recognition test

To evaluate object recognition memory, we introduced a single mouse in an L-maze (15-cm high × 35-cm long × 5-cm wide) during 9 min for three consecutive days (Oliveira da Cruz *et al*, 2020). The first day (habituation session) the maze did not contain any object; the second day (training session) two equal objects (a green object made of Lego pieces) were placed at both ends of the maze; the third day (testing session), a new object, different in shape, colour, and texture (a white and orange object made of Lego pieces) was placed at one of the ends. Position of novel objects in the arms was randomized, and objects were previously analysed not be intrinsically favoured. In all cases, mouse behaviour was video-recorded, and exploration time was manually counted, being exploration considered as mice pointing the nose to the object (distance < 1 cm) whereas biting and standing on the top of the object was not considered exploration. Mice with total exploration times lower than 15 s were considered outliers. Discrimination index was calculated as the time spent exploring the new object (N) minus the time exploring the familiar object (F), divided by the total exploration time [(N-F)/(N+F)]. When administered, SR141716 (Cayman Chemical, Ann Arbor, MI, USA #9000484; 0.3 mg/kg), or vehicle [2% (v/v) DMSO, 2% (v/v) Tween-80 saline solution] was injected intraperitoneally immediately after the training session.

### Fear-conditioning test

To evaluate hippocampal-dependent memory, we conducted a contextual fear-conditioning test. A single mouse was introduced in a fear conditioning chamber (Ugo Basile, Gemonio, VA, Italy #46000) for 2 min, and then 5 electric shocks were applied (0.2 mA for 2 s each, 1-min intervals between shocks). Twenty-four h later, the mouse was reintroduced in the same chamber for 3 min, and freezing behaviour was automatically detected using ANY-maze software (Stoelting Europe, Dublin, Ireland). The latency to start freezing detection was set to two s of immobility. When administered, SR141716 (0.3 mg/kg), or vehicle [2% (v/v) DMSO, 2% (v/v) Tween-80 saline solution] was injected intraperitoneally immediately after the shocking session.

### Y-maze-based memory test

To evaluate hippocampal-dependent memory, we employed a modified version of the Y-maze test (Kraeuter *et al*, 2019). A mouse was placed in one arm of a maze (starting arm) containing three opaque arms orientated at 120° angles from one another, being one arm of the maze closed off (novel arm) and the other open (familiar arm) and allowed for free exploration for 15 min (training session). Position of the starting, familiar, and novel arms was randomized between tests. One h later, the mouse was reintroduced into the maze with all three arms accessible and allowed for free exploration for 5 min (testing session). Animal behaviour was video-recorded, and the total ambulation in each arm was obtained by using Smart3.0 software. In line with equivalent reports (Kraeuter *et al*, 2019), we noted a tendency of the mice to linger at the starting arm, so comparisons were exclusively calculated between the novel arm and the familiar arm. When administered, SR141716 (0.3 mg/kg), or vehicle [2% (v/v) DMSO, 2% (v/v) Tween-80 saline solution)] was injected intraperitoneally the day before the test.

### RNA isolation and quantitative PCR

RNA isolation for multiple tissues was achieved by using the NucleoZOL one phase RNA purification kit (Macherey-Nagel #740404.200) following manufacturer’s instructions. Two µg of total RNA were retro-transcribed using the Transcriptor First Strand cDNA Synthesis Kit (Roche Life Science, Penzberg, Upper Bavaria, Germany, #04379012001) with random hexamer primers. Real-time quantitative RT-PCR (Q-PCR) was performed in a QuantStudio 7/12k Flex System (Applied Biosystems) with the following primers *Crbn*.F 5’-TGAAATGGAAGTTGAAGACCAAGATAG-3; *Crbn*.R 5’-AACTCCTCCATATCAGCTCCCAGG-3’; *Hprt*.F 5’-CAGTACAGCCCCAAAATGGT-3’; *Hprt*.R 5’-CAAGGGCATATCCAACAACA-3’; *Tbp*.F 5’-GGGGAGCTGTGATGTGAAGT-3’; *Tbp*.R 5’-CCAGGAAATAATTCTGGCTCA-3’, using the LightCycler® Multiplex DNA Master (Roche Life Science #07339577001) and SYBR green (Roche Life Science #4913914001). Relative expression ratio was calculated by using the ΔΔCt method with HPRT or TBP as housekeeping genes for normalization.

### RNAscope and immunofluorescence

For RNAscope, mice were deeply anesthetized with a mixture of ketamine/xylazine (87.5 mg/kg and 12.5 mg/kg, of each drug, respectively) and immediately perfused intracardially with PBS followed by 4% paraformaldehyde (Panreac, Barcelona, Spain #252931.1211). After perfusion, brains were removed and post-fixed overnight in the same solution, cryoprotected by immersion in 10, 20, 30% gradient sucrose (24 h for each sucrose gradient) at 4 °C, and then embedded in OCT. Serial coronal cryostat sections (15 μm-thick) through the whole brain were collected in microscope glass slides (Thermo Fisher Scientific, Waltham, MA, USA #J1800AMNZ) and stored at -80 °C. RNAscope assay (Advanced Cell Diagnostics, Newark, California, USA) was performed using RNAscope® Intro Pack for Multiplex Fluorescent Reagent Kit v2 (#323136) with the Crbn mouse probe (#894791) following the manufacturer’s instructions.

For immunofluorescence, serial coronal cryostat sections (30 μm-thick) through the whole brain were collected in PBS as free-floating sections and stored at -20 °C. Slices or coverslips were permeabilized and blocked in PBS containing 0.25% Triton X-100 and 10% or 5% goat serum (Pierce Biotechnology, Rockford, IL, USA), respectively, for 1 h at RT. Primary antibodies were diluted directly into the blocking buffer, and incubated overnight at 4 °C with the following primary antibodies and dilutions: anti-CB_1_R (1:400, CB_1_R-GP-Af530, Frontier Institute Ishikari, Hokkaido, Japan). After 3 washes with PBS for 10 min, samples were subsequently incubated for 2 h at RT with the appropriate highly cross-adsorbed anti-guinea pig AlexaFluor 546, secondary antibody (1:1000; Invitrogen), together with DAPI (Roche, Basel, Switzerland) to visualize nuclei. After washing 3 times in PBS, sections were mounted onto microscope slides using Mowiol® mounting media.

Hybridization and immunofluorescence data were acquired on SP8 confocal microscope (Leica Microsystems, Mannheim, Germany) using LAS-X software. Images were taken using apochromatic 20X objective, and a 3-Airy disc pinhole. Fluorescent quantification was measured using FIJI ImageJ open-source software, establishing a threshold to measure only specific signal that was kept constant along the different images. Regions of interest (ROIs) were defined for CA1 and CA3 pyramidal layer, hilus and granule cell layer of dentate gyrus. Data were then expressed as percentage of control. Controls were included to ensure none of the secondary antibodies produced any significant signal in preparations incubated in the absence of the corresponding primary antibodies. Representative images for each condition were prepared for figure presentation by applying brightness, contrast, and other adjustments uniformly.

### Protein expression and purification

E. coli BL21 DE3 containing pBH4 (pET23-custom derivative) plasmids encoding 6xHis-tagged hCRBN or CB_1_R-CTD (amino acids 400-472) were inoculated in 2 L of 2xYT media (1.6 % w/v tryptone, 1 % w/v yeast extract, and 5 g/L NaCl, pH 7.0) at 37 °C and constant agitation. During the exponential growth phase (OD_600_ = 0.6-0.8), protein expression was induced by addition of 0.5 mM isopropyl 1-thio-β-D-galactopyranoside (Panreac, Barcelona, Spain) for 16 h at 20 °C. Next, bacteria were pelleted by centrifugation at 5,000g for 15 min at room temperature and resuspended in ice-cold lysis buffer (100 mM Tris-HCl, 100 mM NaCl, 10 mM imidazole, pH 7.0) with continuous shaking in the presence of protease inhibitors (1 mg/mL aprotinin, 1 mg/ mL leupeptin, 200 mM PMSF), 0.2 g/L lysozyme, and 5 mM β-mercaptoethanol, followed by four cycles of sonication on ice. Insoluble cellular material was sedimented by centrifugation at 12,000g for 30 min at 4° C and the resultant lysate filtered through porous paper. Recombinant 6xHis-tagged proteins were sequentially purified on a nickel nitrilotriacetic acid affinity column. After extensive washing (50 mM Tris-HCl, 100 mM NaCl, 25 mM imidazole, pH 7.0), proteins were eluted with elution buffer (50 mM Tris-HCl, 100 mM NaCl, 250 mM imidazole, pH 7.0, supplemented with the aforementioned protease inhibitors). Protein purity was confirmed by SDS-PAGE and Coomassie brilliant blue or silver staining. Pure protein solutions were concentrated by centrifugation in Centricon tubes (Millipore).

### Fluorescence polarization

6xHis-tagged CB_1_R-CTD was labelled with 3 molar equivalents of 5-(iodoacetamido)fluorescein (5-IAF) in sodium bicarbonate buffer, pH 9.0, for 1 h at 25

°C, protected from light. Subsequently, non-reacted 5-IAF was washed out with a 1.00-Da cutoff dialysis membrane. The concentration of the labelled peptide was calculated by using the value of 68,000 cm^-1^ M^-1^ as the molar extinction coefficient of the dye at pH 8.0, and a wavelength of 494 nm. Saturation binding experiments were performed essentially as described previously (Costas-Insua *et al*, 2021), with a constant concentration of 100 nM 5-IAF-CB_1_R-CTD and increasing amounts of CRBN (∼0-100 µM), and 3 internal replicates per point within each experiment. The fluorescence polarization values obtained were fitted to the equation (FP – FP0) = (FPmax - FP0)[CRBN]/(Kd + [CRBN]), where FP is the measured fluorescence polarization, FPmax the maximal fluorescence polarization value, FP0 the fluorescence polarization in the absence of added CRBN, and Kd the dissociation constant, as determined with GraphPad Prism version 8.0.1 (GraphPad Software, San Diego, CA, USA).

### Proximity ligation assay (PLA)

*In situ* PLA for CB_1_R and CRBN was conducted in HEK-293T cells transfected with pcDNA3.1-CB_1_R-myc and pcDNA3.1-3xHA-CRBN. Controls were performed in the absence of one of the plasmids, that was replaced by an empty vector. Cells were grown on glass coverslips and fixed in 4 % PFA for 15 min. For conducting PLA in mouse hippocampal brain slices, mice were deeply anesthetized and immediately perfused transcardially with PBS followed by 4 % PFA, postfixed and cryo-sectioned. Immediately before the assay, mouse brain sections were mounted on glass slides, and washed in PBS. In all cases, complexes were detected using the Duolink in situ PLA Detection Kit (Sigma Aldrich) following supplier’s instructions. First, samples were permeabilized in PBS supplemented with 20 mM glycine and 0.05% Triton X-100 for 5 min (cell cultures) or 10 min (mounted slices) at room temperature. Slices were next incubated with Blocking Solution (one drop per cm^2^) in a pre-heated humidity chamber for 1 h at 37 °C. Primary antibodies were diluted in the Antibody Diluent Reagent from the kit [mouse anti-c-myc (clone 9E10; 1:200, Sigma-Aldrich #11667149001) and rabbit anti-HA (1:200, CST, #3724) for cell cultures; rabbit anti-CRBN (1:100, CST, #71810) and rabbit anti-CB_1_R (1:100, Frontier Institute, #CB1-Rb-Af380) for brain sections], and incubated overnight at 4 °C. Negative controls were performed with only one primary antibody. Ligations and amplifications were performed with In Situ Detection Reagent Red (Sigma Aldrich), stained for DAPI, and mounted. Samples were analyzed with a Leica SP8 confocal microscope and processed with Fiji ImageJ software.

### Cannabinoid administration

Adult mice (2–4-month-old) were injected intraperitoneally with vehicle (1% v/v DMSO in 1:18 v/v Tween-80/saline solution) 3 or 10 mg/kg THC (THC Pharm). Forty min later, for the catalepsy test, the animal was placed with both forelimbs leaning on a bar situated at a height of 3.5 cm. Immobility was considered maximal when the animal exceeded 60 s of immobility, and null when the immobility time was lower than 5 s. In all cases, 3 attempts were performed, and the maximal immobility time was selected as the representative value. Next, analgesia was assessed as the latency to paw licking in the hot-plate paradigm at a constant temperature of 52 °C. Animals were assigned randomly to the different treatment groups, and all experiments were performed in a blinded manner for genotype and pharmacological treatment.

### Western blot and immunoprecipitation

Samples for western blotting were prepared as described (Costas-Insua *et al*, 2021; Maroto *et al*, 2023). Tissue samples were homogenized with the aid of an automated grinder (DWK Life Sciences GmbH, Mainz, Germany, #749540-0000). Proteins (1-50 µg) were resolved using PAGE-SDS followed by transfer to PVDF membranes using Bio-Rad FastCast® reagents and guidelines. Membranes were blocked with 5% defatted milk (w/v) or 5% BSA (w/v) in TBS-Tween-20 (0.1%) for 1 h and incubated overnight with the following antibodies and dilutions: anti-phospho-ERK1/2 (1:1,000, CST, Danvers, MA, 333 USA #9101), anti-ERK1/2 (1:1,000, CST #4696), anti-GFP (1:1000, Thermo Fisher Scientific, Waltham, MA, USA #MA5-15256), anti-α-tubulin (1:10,000, Sigma-Aldrich #T9026), anti-β-actin (1:10,000, Sigma-Aldrich #A5441), anti-FLAG M2 (1:1,000, Sigma-Aldrich #F3165), anti-HA (1:1,000, CST #3724), anti-GAPDH (1:3,000, CST #2118), anti-HSP90 (1:3,000 SCBT #sc-69703), anti-CB_1_R (1:2000, CB_1_R-GP-Af530, Frontier Institute Ishikari, Hokkaido, Japan), anti-CRBN (1:1,000, CST #71810), anti-Ubiquitin (SCBT, sc-8017), anti-synaptophysin (Synaptic Systems, Goettingen, Germany #101002), anti-vGLUT1 (Synaptic Systems, #135303), anti-vGAT (Synaptic Systems, #131003), anti-PSD-95 (Abcam, Cambridge, UK, #ab2723), anti-vinculin (1:5,000, Sigma-Aldrich, #V9264). All antibodies were prepared in TBS Tween-20 (0.1%) with 5% BSA (w/v). Membranes were then washed three times with TBS-Tween-20 (0.1%), and HRP-labelled secondary antibodies, selected according to the species of origin of the primary antibodies (Sigma-Aldrich #NA-931 and #NA-934 and Invitrogen #A18769), were added for 1 h at a 1:5,000 dilution in TBS-Tween-20 (0.1%) at room temperature. Finally, protein bands were detected by incubation with an enhanced chemiluminescence reagent (Bio-Rad #1705061). All results provided represent the densitometric analysis, performed with Image Lab software (Bio-Rad), of the band density from the protein of interest vs. the corresponding band density from the loading control. For immunoprecipitations, the pulled-down protein was considered the corresponding loading control. Western blot images were cropped for clarity. Electrophoretic migration of molecular weight markers is depicted on the left-hand side of each blot.

Immunoprecipitation experiments were performed as previously (Costas-Insua *et al*, 2021). For co-immunoprecipitation experiments in HEK-293T cells, samples were prepared on ice-cold GST buffer (50 mM Tris-HCl, 10% glycerol v/v, 100 mM NaCl, 2 mM MgCl2, 1% v/v NP-40, pH 7.4), supplemented with protease inhibitors. Denaturing immunoprecipitation to detect ubiquitination was conducted on RIPA buffer (50 mM Tris-HCl pH 7.4, 150 mM NaCl, 1% v/v NP-40, 0.5% w/v sodium deoxycholate, 0.1% w/v sodium dodecyl sulfate) supplemented with the deubiquitinase inhibitor 2-chloroacetamide. Immunoprecipitations were conducted with anti-FLAG M2 affinity gel (Sigma-Aldrich #A2220) or anti-HA agarose (Thermo Scientific, #26181), following the supplier instructions. Finally, for co-immunoprecipitation experiments in adult hippocampal tissue, protein extracts were solubilized on DDM buffer (25 mM Tris-HCl pH 7.4, 140 mM NaCl, 2 mM EDTA, 0.5% n-dodecyl-β-D-maltoside) and the following antibodies were added to a final concentration of 1 µg/ml: anti-CRBN (CST #71810), anti-CB_1_R (CB_1_R-Rb-Af380), IgG control (Thermo Fisher Scientific, #10500C). Bound proteins were captured with Protein G agarose for 4 h (Sigma-Aldrich, #17061801), spun at low speed, washed three times with lysis buffer, and eluted with 2x Laemmli sample buffer. In all cases, for CB_1_R immunodetection, samples were heated for 10 min at 55 °C, and appropriate CB_1_R-KO controls were included, following recommended guidelines (Esteban *et al*, 2020).

### Cell culture, transfection and signalling experiments

The HEK-293T cell line was obtained from the American Type Culture Collection (Manassas, VA, USA). HEK-293T-*CRBN-KO* and parental HEK-293T-*CRBN-WT* cells, generated with CRISPR/Cas9 technology, (Krönke *et al*, 2015), were kindly provided by Dr. Benjamin L. Ebert (Dana-Farber Cancer Institute, Boston, MA, USA). Cells were grown in DMEM supplemented with 10% FBS (Thermo Fisher Scientific), 1% penicillin/streptomycin, 1 mM Na-pyruvate, 1 mM L-glutamine, and essential medium non-essential amino acids solution (diluted 1/100) (all from Invitrogen, Carlsbad, CA, USA). Cells were maintained at 37 °C in an atmosphere with 5% CO_2_, in the presence of the selection antibiotic when required (HEK-293T-FLAG-CB_1_R; zeocin at 0.22 mg/mL, Thermo Fisher Scientific #R25001), and were periodically checked for the absence of mycoplasma contamination. Cell transfections were conducted with polyethyleneimine (Polysciences inc. Warrington, PA, USA #23966) in a 4:1 mass ratio to DNA according to the manufacturer’s instructions. Double transfections were performed with equal amounts of the two plasmids (5 µg of total DNA per 10-cm plate), except for BRET experiments (see below). Every condition was assayed in triplicate within each individual experiment.

Drug treatments to assess CB_1_R-evoked signalling were conducted as follows. For ERK phosphorylation experiments, a 10 cm-diameter plate of transfected cells was trypsinized and seeded on different 6 cm-diameter plates at a density of 1×10^6^ cells per well. Six h later, cells were serum-starved overnight. Then, WIN-55,212-2 (Sigma-Aldrich; #W102, 0.01-1 µM final concentration) or vehicle (DMSO, 0.1% v/v final concentration) was added for 10 min. For PKA activity assays, the procedure was essentially the same, but following WIN-55,212-2 (1 µM final concentration) or vehicle (DMSO, 0.1% v/v final concentration) treatment, forskolin (Tocris, Bristol, UK, #1099, 1 µM final concentration) or vehicle (DMSO, 0.1% v/v final concentration) was added for another 10 min. Cells were subsequently washed with ice-cold PBS, snap-frozen in liquid nitrogen, and harvested at -80 ° C for western blot analyses, except for the determination of PKA activity by ELISA (see below). Every condition was assayed in triplicate within each individual experiment.

### Bioluminescence resonance energy transfer (BRET)

BRET was conducted as described (Costas-Insua *et al*, 2021) in HEK-293T cells transiently co-transfected with a constant amount of cDNA encoding the receptor fused to Rluc protein and with increasingly amounts of GFP-CRBN. The net BRET is defined as [(long-wavelength emission)/(short-wavelength emission)] – Cf where Cf corresponds to [(long-wavelength emission)/(short-wavelength emission)] for the Rluc construct expressed alone in the same experiment. BRET is expressed as milli BRET units (mBU; net BRET x 1000). In BRET curves, BRET was expressed as a function of the ratio between fluorescence and luminescence (GFP/Rluc). To calculate maximal BRET from saturation curves, data were fitted using a nonlinear regression equation and assuming a single phase with GraphPad Prism software version 8.0.1. The represented experiment is the mean of three biological replicates.

### Antibody-capture [^35^S]GTPγS scintillation proximity assay

CB_1_R-mediated activation of different subtypes of Gα protein subunits (Gα_i1_, Gα_i2_, Gα_i3_, Gα_o_, Gα_q/11_, Gα_s_, Gα_z_, and Gα_12/13_) was determined as described (Costas-Insua *et al*, 2021) using a homogeneous protocol of [^35^S]GTPγS scintillation proximity assay coupled to the use of the following antibodies: mouse monoclonal anti-Gα_i1_ (1:20, Santa Cruz Biotechnology #sc-13534), rabbit polyclonal anti-Gα_i2_ (1:20; Santa Cruz Biotechnology #sc-7276), rabbit polyclonal anti-Gα_i3_ (1:60, Antibodies on-line #ABIN6258933), mouse monoclonal anti-Gα_o_ (1:40, Santa Cruz Biotechnology #sc-393874), mouse monoclonal anti-Gα_q/11_ (1:20, Santa Cruz Biotechnology #sc-515689), rabbit polyclonal anti-Gα_s_ (1:20, Santa Cruz Biotechnology #sc-377435), rabbit polyclonal anti-Gα_z_ (1:60, Antibodies on-line #ABIN653561), and rabbit polyclonal anti-Gα_12/13_ (1:40, Antibodies on-line #ABIN2848694). To determine their effect on [^35^S]GTPγS binding to the different Gα subunit subtypes in the different experimental conditions, a single submaximal concentration (10 µM) of WIN-55,212-2 or HU-210 (Tocris #0966) was used, either alone or in the presence of the CB_1_R antagonist O-2050 (10 µM, Tocris #1655) as control. Nonspecific binding was defined as the remaining [^35^S]GTPγS binding in the presence of 10 µM unlabelled GTPγS. For each Gα protein, specific [^35^S]GTPγS binding values were transformed to percentages of basal [^35^S]GTPγS binding values (those obtained in the presence of vehicle). Every condition was assayed in triplicate within each individual experiment.

### Determination of cAMP concentration

cAMP was determined using the Lance Ultra cAMP kit (PerkinElmer), which is based on homogeneous time-resolved fluorescence energy transfer. Briefly, HEK-293T cells (1,000 per well), growing in medium containing 50 µM zardeverine, were incubated for 15 min in white ProxiPlate 384-well microplates (PerkinElmer) at 25 °C with vehicle WIN-55,212-2 or CP55,940 (doses ranging from 0.0025 to 1 µM final concentration) before adding vehicle or forskolin (0.5 μM final concentration) and incubating for 15 additional min. Every condition was assayed in triplicate within each individual experiment. Fluorescence at 665 nm was analysed on a PHERAstar Flagship microplate reader equipped with an HTRF optical module (BMG Lab technologies, Offenburg, Germany).

### Dynamic mass redistribution (DMR) assays

Global CB_1_R signalling was determined by label-free technology as previously described (Costas-Insua *et al*, 2021; Maroto *et al*, 2023) by using an EnSpire® Multimode Plate Reader (PerkinElmer, Waltham, MA, USA). Briefly, 10,000 HEK-293T or HEK-293T-Crbn^-/-^ cells expressing CB_1_R were plated in 384-well sensor microplates and cultured for 24 h. Then, the sensor plate was scanned, and a baseline optical signature was recorded before adding 10 μL of the cannabinoid receptor agonist WIN-55,212-2 (Sigma-Aldrich, 100 nM final concentration) dissolved in assay buffer (HBSS with 20 mM Hepes, pH 7.15) containing 0.1% DMSO. Then, the resulting shifts of reflected light wavelength (in pm) were analysed by using EnSpire Workstation Software version 4.10. Each representative curve shown is the mean of three different experiments. When conducted, cell transfection was achieved as stated above.

### Plasmids

3xFLAG-tagged human CB_1_R was cloned in the pcDNA3.1 backbone by restriction cloning from existing sources in our laboratory. *N*-terminal 3xHA-tagged cDNAs of mouse and human CRBN, as well as V5-tagged human Cullin-4a and myc-tagged human DDB1 were acquired to VectorBuilder (Chicago, IL, USA). The GFP-tagged version, partial and deletion mutants of CRBN were built by conventional PCR methods. His_6_-tagged CB_1_R-CTD, CB_1_R-CTD mutants, CB_1_R-myc and CB_1_R-Rluc were already made in a previous work (Costas-Insua *et al*, 2021). Human CRBN cDNA was inserted in the pBH4 vector by restriction cloning, rendering a His_6_-tagged CRBN amenable for protein purification; or in the pAM-CBA (Ruiz-Calvo *et al*, 2018) plasmid for adeno-associated viral particles production (see below).

### Adeno-associated viral vector production

All vectors used were of an AAV1/AAV2 mixed serotype and were generated by calcium phosphate transfection of HEK293T cells. Subsequent purification was conducted using an iodixanol gradient and ultracentrifugation as described previously (Maroto et al, 2023).

### Stereotaxic surgery

Adult mice (2 months-old) were anaesthetized with isoflurane (4%) and placed into a stereotaxic apparatus (World Precision Instruments, Sarasota, FL, US). Adeno-associated viral particles were injected with a Hamilton microsyringe (Sigma-Aldrich #HAM7635-01) coupled to a 30g-needle controlled by a pump (World Precision Instruments, #SYS-Micro4) directly in the hippocampus (1 µL per injection site at a rate of 0.25 µL/min) with the following coordinates (in mm): anterior-posterior: -2.00 mm, dorsal-ventral: -2.00 and -1.5 mm, medial-lateral: ±1.5 mm. Following each injection, the syringe remained positioned for 1 min before withdrawal. Mice were treated with analgesics [buprenorphine (0.1 mg/kg) and meloxicam (1 mg/kg)] before and for three consecutive days after surgery. After three weeks of recovery, once ensured that body weight returned at least to pre-surgery values, mice were euthanized, and brain was dissected to collect hippocampi for further procedures.

### Determination of PKA activity

To determine CB_1_R-induced inhibition of PKA, we employed an ELISA (Abcam, ab139435) following the manufacturer’s instructions. Briefly, HEK-293T cells stably expressing CB_1_R, treated or not with WIN55,212-2 and/or forskolin, as stated above, were lysed immediately after treatment with assay buffer (20 mM MOPS, 50 mM β-glycerophosphate, 50 mM sodium fluoride, 1 mM sodium orthovanadate, 5 mM EGTA, 2 mM EDTA, 1% NP40, 1 mM DTT, 1 mM benzamidine, 1 mM PMSF, 10 µg/mL leupeptin and aprotinin). The amount of total protein assayed (1-50 µg) was independently adjusted in each assay, to ensure a linear protein-signal dependency. A positive control, consisting of increasing amounts of recombinant PKA, was included in each independent experiment. Every condition was assayed in triplicate within each individual experiment.

### CRBN knockdown

Silencing of CRBN was achieved by transfecting HEK-293T cells with the following stealth siRNAs (Invitrogen) (Ito *et al*, 2010) using Lipofectamine 2000 (Thermo Fisher Scientific #11668027) according to the manufacturer’s instructions: CRBN #1, 5’-CAGCUUAUGUGAAUCCUCAUGGAUA-3’; CRBN #2, 5’-CCCAGACACUGAAGAUGAAAUAAGU-3’. Only sense strands are shown. Stealth RNAi of low GC content was included as a negative control.

### Experimental design and statistical analyses

Unless otherwise indicated, data are presented as mean ± SEM. The particular statistical tests that were applied are indicated in each figure legend. All datasets were tested for normality and homoscedasticity prior to analysis. Whenever possible, the precise p values are given in the figures. p values below 0.05 were considered significant. The sample size for each experiment was estimated based on previous studies conducted by our laboratories. The number of biological replicates is provided in each figure legend. The number of technical replicates is provided in the corresponding Materials and Methods subsection. Graphs and statistics were generated by GraphPad Prism v8.0.1.

## Acknowledgements

This work was supported by the Spanish *Ministerio de Ciencia e Innovación* (MICINN/FEDER; grants PID2021-125118OB-I00 to M.G., PID2020-113938RB-I00 to E.M. and V.C., and PID2019-106404RB-I00 to L.U.,) and by the *Generalitat de Catalunya* (grant 2021-SGR-00230 to E.M. and V.C.) L.B. was supported by INSERM. C.C.-I. and I.B.M. were supported by contracts from the Spanish *Ministerio de Universidades* (*Formación de Profesorado Universitario* Program, references FPU16/02593 and FPU15/01833, respectively). We are indebted to Dr. Benjamin L. Ebert for the kind donation of HEK-293T-*CRBN-KO* and parental HEK-293T-*CRBN-WT* cells. We also thank David Martín-Gutiérrez, Lucía Rivera-Endrinal, Dr. Daniel García-Ovejero, Dr. Eduardo Molina-Holgado, and the personnel of the core microscopy centre and the animal facilities of Complutense University of Madrid for their expert technical assistance.

## Author contributions

**Carlos Costas-Insua:** Conceptualization; data curation; formal analysis; investigation; methodology; resources; software; supervision; validation; visualization; writing – original draft; writing – review & editing. **Alba Hermoso-López:** Data curation; formal analysis; investigation; methodology; software; validation; visualization; writing – review & editing. **Estefanía Moreno:** Data curation; formal analysis; investigation; methodology; resources; software; validation; visualization; writing – review & editing. **Carlos Montero-Fernández:** Investigation; methodology; software; writing – review & editing. **Alicia Álvaro-Blázquez:** Investigation; methodology; software; writing – review & editing. **Rebeca Diez-Alarcia:** Formal analysis; investigation; methodology; software; writing – review & editing. **Irene B. Maroto:** Formal analysis; investigation; methodology; software; validation; visualization; writing – review & editing. **Paula Morales:** Data curation; formal analysis; investigation; methodology; resources; software; visualization; writing – review & editing. **Enric I. Canela:** Funding acquisition; methodology; resources; supervision; writing – review & editing. **Vicent Casadó:** Funding acquisition; methodology; resources; supervision; writing – review & editing. **Leyre Urigüen:** Data curation; formal analysis; funding acquisition; methodology; resources; software; supervision; validation; writing – review & editing. **Luigi Bellocchio:** Formal analysis; funding acquisition; methodology; resources; supervision; writing – review & editing. **Ignacio Rodríguez-Crespo:** Conceptualization; data curation; formal analysis; methodology; resources; software; supervision; validation; writing – review & editing. **Manuel Guzmán:** Conceptualization; data curation; formal analysis; funding acquisition; methodology; project administration; supervision; validation; visualization; writing – original draft; writing – review & editing.

## The Paper Explained

### Problem

Intellectual disability is a major healthcare problem. Specifically, disrupting mutations in *CRBN*, the gene that encodes cereblon/CRBN, an E3 ubiquitin ligase complex component, cause a form of autosomal recessive non-syndromic intellectual disability (ARNSID) that heavily impairs learning and memory skills. Recently, owing to the generation of *Crbn* knockout mice that recapitulate the human disease, some molecular factors underlying that cognitive dysfunction have been proposed, but the intimate CRBN deficiency-evoked etiopathological mechanisms remain unknown.

### Results

We first developed mouse models in which the *Crbn* gene was knocked-out non-selectively from all body cells (CRBN-KO), or selectively from the glutamatergic (Glu-CRBN-KO) or GABAergic (GABA-CRBN-KO) forebrain-neuron lineage. Behavioural testing revealed a profound memory impairment in CRBN-KO and Glu-CRBN-KO but not CRBN-GABA-KO mice. Molecular studies demonstrated that CRBN interacts physically with CB_1_R and inhibits receptor action in a ubiquitin ligase-independent manner, thus providing a rationale for the CB_1_R overactivation displayed by CRBN-deficient animals. Finally, experiments conducted with CRBN-KO and Glu-CRBN-KO mice acutely treated with rimonabant, a CB_1_R-selective antagonist, showed that blockade of this receptor restores normal memory function.

### Impact

Our findings demonstrate that *i*) CRBN binds to and inhibits CB_1_R, *ii*) deleting CRBN causes CB_1_R overactivation, and *iii*) this event, in turn, drives CRBN deficiency-associated memory deficits in mice. In full caption, our findings pave the way for the pharmacological blockade of CB_1_R as a novel therapeutic intervention in patients with CRBN deficiency-linked ARNSID.

## Data Availability Section

This study includes no data deposited in external repositories.

## EXPANDED VIEW FIGURE LEGENDS

**Figure EV1.**
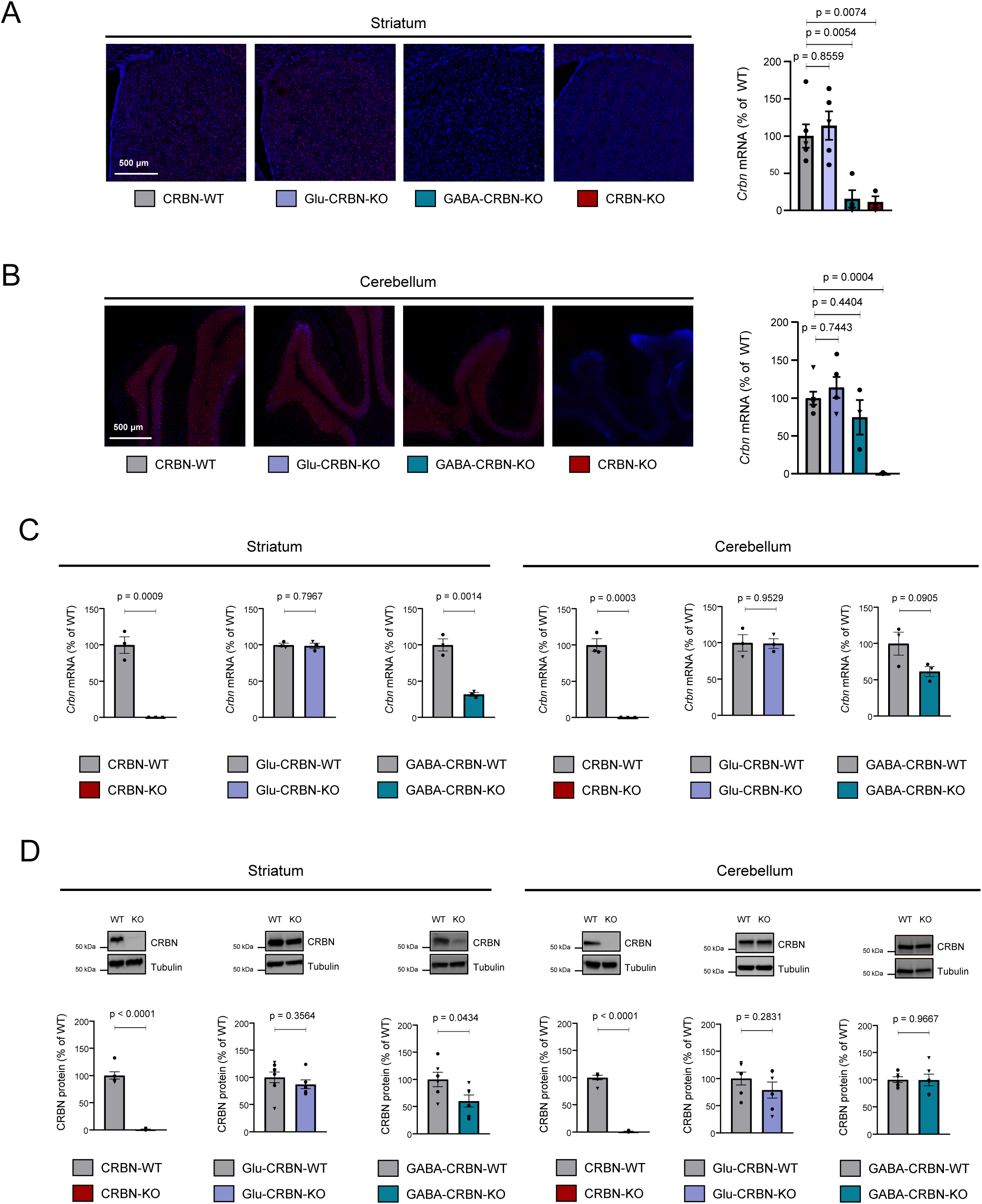
Additional characterization of the conditional CRBN knockout mouse lines. A. Representative images and fluorescent signal quantification of RNAscope *in situ* hybridization labelling in the striatum of CRBN-WT (n = 6), Glu-CRBN-KO (n = 5), GABA-CRBN-KO (n = 4) and CRBN-KO (n = 3) mice. Circles, male mice; triangles, female mice. p values were obtained by one-way ANOVA with Dunnett’s post-hoc test. B. Representative images and fluorescent signal quantification of RNAscope *in situ* hybridization labelling in the cerebellum of CRBN-WT (n = 6), Glu-CRBN-KO (n = 5), GABA-CRBN-KO (n = 3) and CRBN-KO (n = 3) mice. Circles, male mice; triangles, female mice. p values were obtained by one-way ANOVA with Dunnett’s post-hoc test. C. *Crbn* mRNA levels (% of WT mice) as assessed by q-PCR in the striatum or cerebellum of CRBN-WT, CRBN-KO, Glu-CRBN-WT, Glu-CRBN-KO, GABA-CRBN-WT and GABA-CRBN-KO mice (n = 3 animals per group). Circles, male mice; triangles, female mice. p values were obtained by unpaired Student’s *t* test. D. CRBN protein levels (% of WT mice) as assessed by western blotting in the striatum or cerebellum of CRBN-WT, CRBN-KO, Glu-CRBN-WT, Glu-CRBN-KO, GABA-CRBN-WT and GABA-CRBN-KO mice (n = 6 animals per group). Circles, male mice; triangles, female mice. p values were obtained by unpaired Student’s *t* test.

**Figure EV2.**
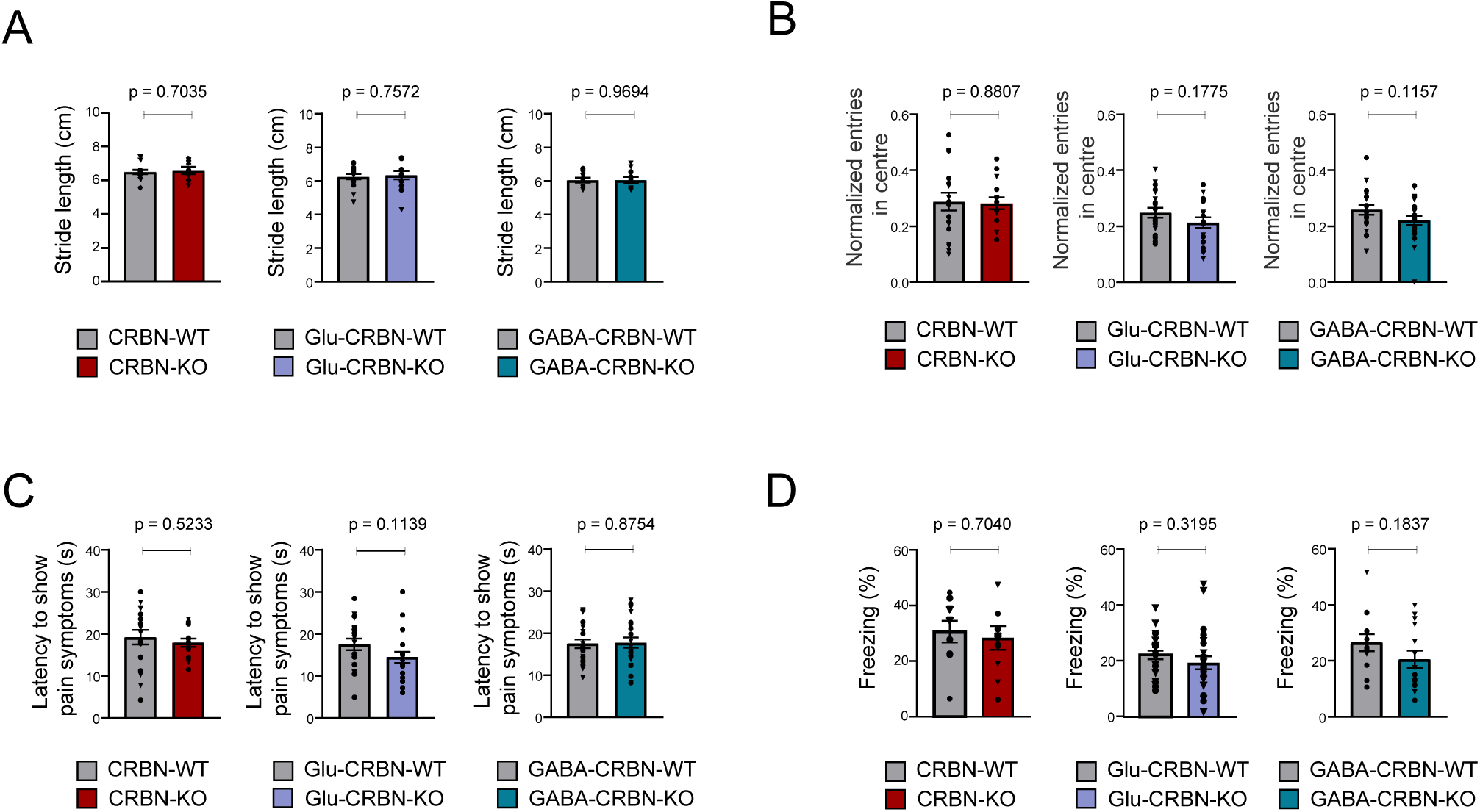
Additional behavioural phenotyping of the CRBN knockout mouse lines. A. Stride length (in cm) in the footprint test. CRBN-WT (n = 13), CRBN-KO (n = 8), Glu-CRBN-WT (n = 17), Glu-CRBN-KO (n = 13), GABA-CRBN-WT (n = 9), GABA-CRBN-KO (n = 11). Circles, male mice; triangles, female mice. p values were obtained by unpaired Student’s *t* test. B. Entries in the central part of an open field arena (normalized to total ambulation). CRBN-WT (n = 18), CRBN-KO (n = 15), Glu-CRBN-WT (n = 20), Glu-CRBN-KO (n = 19), GABA-CRBN-WT (n = 19), GABA-CRBN-KO (n = 24). Circles, male mice; triangles, female mice. p values were obtained by unpaired Student’s *t* test. C. Time to show pain symptoms (in s) in the hot plate test. CRBN-WT (n = 18), CRBN-WT (n = 18), CRBN-KO (n = 15), Glu-CRBN-WT (n = 20), Glu-CRBN-KO (n = 19), GABA-CRBN-WT (n = 21), GABA-CRBN-KO (n = 24). Circles, male mice; triangles, female mice. p values were obtained by unpaired Student’s *t* test. D. Time (in %) spent freezing in the conditioning session of the fear conditioning protocol. CRBN-WT (n = 10), CRBN-KO (n = 10), Glu-CRBN-WT (n = 24), Glu-CRBN-KO (n = 24), GABA-CRBN-WT (n = 13), GABA-CRBN-KO (n = 14). Circles, male mice; triangles, female mice. p values were obtained by unpaired Student’s *t* test.

**Figure EV3.**
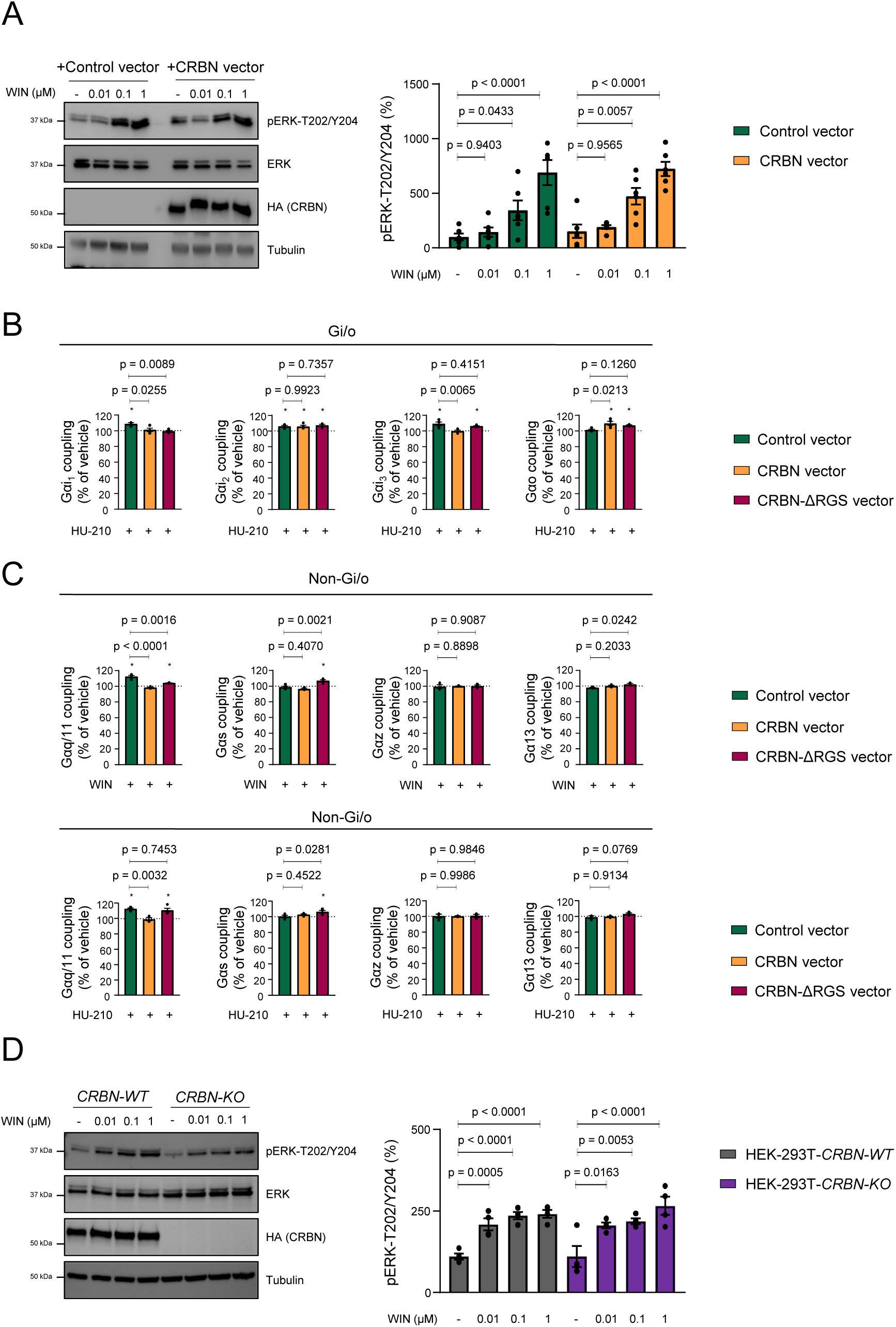
Additional data on the CRBN-mediated inhibition of CB_1_R-evoked G_i/o_ protein signalling *in vitro*. A. HEK-293T cells expressing CB1R, together or not with CRBN, were incubated for 10 min with vehicle or WIN55,212-2 (doses ranging from 0.01 to 1 µM), and cell extracts were blotted for ERK phosphorylation. A representative experiment is shown. p values were obtained by two-way ANOVA with Tukey’s multiple comparisons test (n = 6). B. Coupling of CB_1_R to Gα_i/o_ proteins in membrane extracts from HEK-293T cells expressing CB_1_R, together or not with CRBN or CRBN-ΔRGS, after HU-210 stimulation (10 µM). *p<0.05 from basal (dashed line) by one-sample Student’s t-test. p values between constructs were obtained by unpaired Student’s *t* test (n = 3-4). C. Coupling of CB_1_R to non-Gα_i/o_ proteins in membrane extracts from HEK-293T cells expressing CB_1_R, together or not with CRBN or CRBN-ΔRGS after WIN55,212-2 stimulation (10 µM). *p<0.05 from basal (dashed line) by one-sample Student’s *t* test. p values between constructs were obtained by unpaired Student’s *t* test (n = 3-4). D. HEK-293T-*CRBN-WT* and HEK-293T-*CRBN-KO* cells expressing CB_1_R were incubated for 10 min with vehicle or WIN55,212-2 (doses ranging from 0.01 to 1 µM), and cell extracts were blotted for ERK phosphorylation. A representative experiment is shown. p values were obtained by two-way ANOVA with Tukey’s multiple comparisons test (n = 4).

**Figure EV4.**
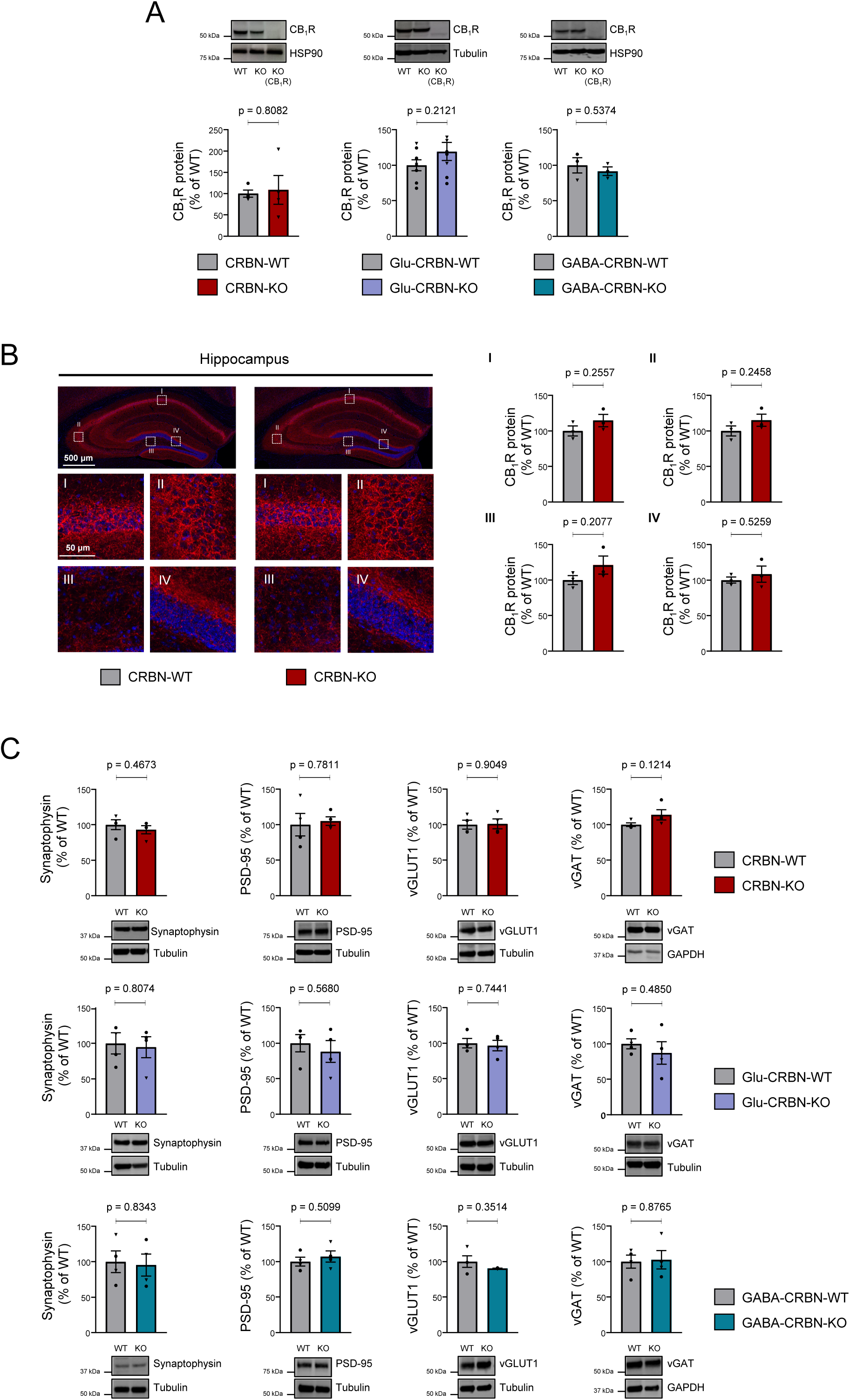
*Crbn* deletion does not alter the levels of CB_1_R and synapse-marker proteins in the mouse hippocampus. A. CB_1_R protein levels (% of WT mice) as assessed by western blotting in the hippocampus of CRBN-WT (n = 4), CRBN-KO (n = 4), Glu-CRBN-WT (n = 8), Glu-CRBN-KO (n = 8), GABA-CRBN-WT (n = 3) or GABA-CRBN-KO (n = 3) mice. Circles, male mice; triangles, female mice. p values were obtained by unpaired Student’s *t* test. B. CB_1_R immunoreactivity (% of WT mice) in the hippocampus of CRBN-WT and CRBN-KO mice (n = 3 animals per group). High magnification images of CA1 (I), CA3 (II), hilus (III) and granule cell layer of the dentate gyrus (IV) are shown. Circles, male mice; triangles, female mice. p values were obtained by unpaired Student’s *t* test. C. Synaptophysin, PSD-95, vGLUT1 and vGAT protein levels (% of WT mice) as assessed by western blotting in the hippocampus of CRBN-WT, CRBN-KO, Glu-CRBN-WT, Glu-CRBN-KO, GABA-CRBN-WT or GABA-CRBN-KO mice (n = 3-4 animals per group). Circles, male mice; triangles, female mice. p values were obtained by unpaired Student’s *t* test

